# Neurocomputational mechanisms underlying emotional awareness: insights afforded by deep active inference and their potential clinical relevance

**DOI:** 10.1101/681288

**Authors:** Ryan Smith, Richard D. Lane, Thomas Parr, Karl J. Friston

**Author notes:** Corresponding Author Information: Ryan Smith, Laureate Institute for Brain Research, 6655 S Yale Ave, Tulsa, OK 74136, USA.

## Abstract

Emotional awareness (EA) is recognized as clinically relevant to the vulnerability to, and maintenance of, psychiatric disorders. However, the neurocomputational processes that underwrite individual variations remain unclear. In this paper, we describe a deep (active) inference model that reproduces the cognitive-emotional processes and self-report behaviors associated with EA. We then present simulations to illustrate (seven) distinct mechanisms that (either alone or in combination) can produce phenomena – such as somatic misattribution, coarse-grained emotion conceptualization, and constrained reflective capacity – characteristic of low EA. Our simulations suggest that the clinical phenotype of impoverished EA can be reproduced by dissociable computational processes. The possibility that different processes are at work in different individuals suggests that they may benefit from distinct clinical interventions. As active inference makes particular predictions about the underlying neurobiology of such aberrant inference, we also discuss how this type of modelling could be used to design neuroimaging tasks to test predictions and identify which processes operate in different individuals – and provide a principled basis for personalized precision medicine.

## 1. Introduction

Trait emotional awareness (tEA) – the ability to conceptualize and understand one’s own affective responses – is now recognized as an important source of individual variability in both clinical psychology and neuroscience (Barrett, 2017; Lane et al., 2015; Panksepp et al., 2017; Smith et al., 2015; R Smith et al., 2017b, 2017c; Smith et al., 2018d, 2018b, 2018c, 2019e; Smith and Lane, 2015, 2016; Wright et al., 2017). Attempts to operationalize this variability have led to a range of overlapping constructs, including levels of emotional awareness (Lane and Schwartz, 1987), emotion differentiation or granularity (Kashdan et al., 2015; Kashdan and Farmer, 2014), and alexithymia (Bagby et al., 1994a, 1994b).

This aforementioned body of work is largely motivated by the clinical relevance of a person’s ability to understand the emotions of self and others. In previous studies based on the theory of levels of emotional awareness (Lane and Schwartz, 1987), lower awareness levels have been associated with several psychiatric disorders and poorer physical health (Baslet et al., 2009; Berthoz et al., 2000; Bydlowski et al., 2005; Consoli et al., 2010; Donges et al., 2005; Frewen et al., 2008; Lackner, 2005; Levine et al., 1997; Moeller et al., 2014; Subic-Wrana et al., 2005, 2007); higher levels have instead been associated with a range of adaptive emotion-related traits and abilities (Barchard and Hakstian, 2004; Bréjard et al., 2012; Ciarrochi et al., 2003; Lane et al., 2000, 1996, 1990). Multiple evidence-based psychotherapeutic approaches also aim (although some more explicitly than others) to improve emotional awareness as a central part of psycho-education in psychotherapy (Barlow et al., 2016; Burum and Goldfried, 2007; Hayes and Smith, 2005).

Here, we focus on the construct of levels of emotional awareness. Within the theory of levels of emotional awareness (tLEA), and its accompanying measurement scale (the levels of emotional awareness scale), five different categorical levels are distinguished (although these are understood to reflect particular points on a continuum; (Lane et al., 1990; Lane and Schwartz, 1987)). At the lowest two levels, subjective awareness of emotions is restricted to somatic sensations and valenced approach-avoidance tendencies, respectively. That is, a person with “level 1” emotional awareness would tend to somatize, in the sense that they may misattribute emotion-related sensations from their body to physical illness or disease, whereas a person with “level 2” emotional awareness may simply recognize that they feel emotionally “bad” or that they “feel like running away”. The third level corresponds to subjective categorization using distinct emotion categories, such as awareness of sadness, anger, and fear. The fourth level involves the ability to entertain multiple emotions in mind simultaneously, such as feeling a blend of anger and fear. The highest level corresponds to an additional theory of mind ability, where an individual is further able to distinguish the emotions of self and others.

Aside from its clinical relevance described above, a number of neuroimaging studies have also attempted to characterize the neural basis of tEA. Based in part on this work, a “three-process model” (TPM; (Smith et al., 2018a; Smith, 2019; Smith et al., 2019b) has recently been proposed to distinguish a range of processes that contribute to emotion episodes, and how individual differences in these processes could contribute to trait differences in emotional awareness (and in the subsequent use of this awareness to guide adaptive decision-making). The TPM distinguishes the following three broad processes:

1. *Affective response generation*: a process in which somatovisceral and cognitive processes are automatically modulated in response to an affective stimulus (whether real, remembered, or imagined) in a context-dependent manner, based on an (often implicit) appraisal of the salience of that stimulus for the survival and goal-achievement of the individual and of associated upcoming metabolic or behavioral demands.
2. *Affective response representation*: a process in which the somatovisceral component of an affective response is subsequently perceived via afferent sensory processing, and then conceptualized under a particular emotion category (e.g., sadness, anger, etc.) in consideration of all other available sources of information (e.g., stimulus or context information, current thoughts or beliefs about the situation, etc.).
3. *Conscious access*: a process in which somatovisceral percepts and emotion concepts are made accessible to domain-general cognition and can be held in working memory – allowing the use of this information in goal-directed decision-making (e.g., verbal reporting, selection of voluntary emotion regulation strategies, etc.).^1^

While this theory has proposed a tentative mapping between these processes and large-scale brain structure and function, their neuro-computational implementation has received less attention. The computational level of description can provide additional mechanistic insights, which could potentially be leveraged to inform and improve clinical interventions. While previous theoretical work has applied active inference to emotional phenomena (Allen and Friston, 2018; Barrett and Simmons, 2015; Clark et al., 2018; Joffily and Coricelli, 2013; Owens et al., 2018; Seth, 2013; Seth and Friston, 2016; R Smith et al., 2017c, 2017a; Smith et al., 2019a), formal modeling of the processes currently under discussion has only recently begun to emerge (e.g., for formal modeling work with respect to emotion concept learning, see (Smith et al., 2019d)). This type of modeling may be central to addressing a significant issue in research on the development and selection of individualized interventions: the fact that more than one underlying aberrant neural (and/or cognitive) process can produce the same measurable symptoms and clinical signs (Anderson, 2014). In the context of the present discussion, this suggests that two individuals could have equally low levels of emotional awareness as measured by current research instruments – even if distinct underlying mechanisms are responsible for this difference. As such, a particular intervention could target the relevant mechanism for one individual but not for the other. Because computational approaches can characterize underlying processes that generate observable outcomes, they are promising both in highlighting the relevant processes and motivating the development of more sensitive measures that can disambiguate process theories on an individual basis.

In this manuscript, we provide an in-depth theoretical example – using the levels of emotional awareness construct – to illustrate how a computational model can reproduce a clinical phenomenon of interest and afford insights about processes relevant to treatment selection. Specifically, we present a hierarchical (deep temporal) neuro-computational model motivated by the active inference framework (Friston et al., 2017a, 2016) that can account for multiple levels of emotional awareness (EA). This model allows for quantitative simulations of how affective bodily responses can be represented (EA level 1/2) and categorized as emotions (EA level 3), and how multiple emotions – including those of both oneself and others – can then be held in mind to inform verbal reports and goal-directed decisions (EA level 4/5). We do not fully address the theory of mind abilities associated with the fifth level of emotional awareness; however, the current report serves as a foundation for future work along these lines.

We first provide a brief primer on active inference, and then describe our model and the results of a number of simulations. These simulations illustrate the way individual differences in (seven) distinct underlying deep inference processes result in measurable phenotypes associated with different levels of emotional awareness. Our simulations will also show that, based on the neural process theory accompanying active inference (Friston et al., 2017a), predictions about empirically measurable individual differences in neuronal and behavioral responses can, in principle, be generated and used to identify which underlying processing mechanisms are most likely contributing to low emotional awareness in different individuals. After presenting these simulations, we highlight how a better understanding of the plurality of processes could inform the development and selection of clinical interventions on an individualized basis.

## 2. Active inference

According to active inference (Friston et al., 2017a), the brain is an inference machine that approximates probabilistic (Bayesian) belief updating across perceptual, cognitive, and motor domains. From this perspective, the brain embodies a generative model that can simulate (i.e., generate predictions about) the sensory data that it would receive if its model of the world was apt. Predicted sensory data can be compared to sensory inputs, and differences between predicted and observed sensations can be used to update the model. Over short timescales (e.g., a single sensory observation), this updating corresponds to *inference* (perception), whereas on longer timescales it corresponds to *learning* (i.e., updating expectations about how external causes generate patterns of sensory input). In other words, perception optimizes beliefs about the current state of the world, while learning optimizes beliefs about the relationships between states of the world and their implicit contingencies. This can be seen as ensuring the generative model remains an accurate model of the world that it seeks to regulate (Conant and Ashbey, 1970).

Active inference casts decision-making in terms of uncertainty reduction; c.f., planning as inference. Actions are chosen to resolve uncertainty about states of affairs under a generative model (i.e., sampling from domains in which the model does not make precise predictions). This active sampling of sensory outcomes minimizes anticipated deviations from predicted outcomes. Actions can also be chosen to realize the observations that an agent prefers (e.g., observing preferred amounts of warmth, social support, food, etc.). Under active inference, such preferences are also formally treated as expectations. In other words, if the agent “expects” to observe her preferred actions, actions will be chosen so as to fulfill those expectations – thereby minimizing surprise and uncertainty. Based on the formalism underlying active inference (described further below), actions are typically first chosen to minimize uncertainty about the environment; once the agent is confident in her understanding of the environmental context in which she is operating, action then becomes goal-seeking (i.e., to fulfill prior preferences). Mathematically, this can be described as selecting sequences of actions (policies) that maximize “Bayesian model evidence” expected under a policy, where model evidence is the (marginal) likelihood that particular sensory inputs would be observed under a given model, which is characterized by a set of prior preferences.

In real-world settings, directly computing Bayesian model evidence is generally intractable. Thus, some approximation is necessary. Active inference uses an approximation based on minimizing a quantity called “variational free energy”, which provides a bound on model evidence, such that model evidence is maximized indirectly. In this case, decision-making is approximately (Bayes) optimal, if the model infers (and enacts) the policy that will minimize expected free energy (i.e., free energy with respect to a policy, where one takes expected future observations into account).

Expected free energy can be decomposed in different ways that reflect uncertainty and prior preferences, respectively (e.g., epistemic and instrumental affordance or ambiguity and risk). This formulation means that any agent that minimizes expected free energy generally engages in exploratory behavior to minimize uncertainty in a new environment. Once uncertainty is resolved, the agent then uses her acquired familiarity with the environment to choose actions that fulfill prior preferences. The formal basis for active inference has been detailed elsewhere (Friston et al., 2017a), and the interested reader is referred to this previous work for a full mathematical treatment.

When the generative model is formulated as a partially observable Markov decision process – a mathematical model of decision-making in cases where states of the world are not directly known but must be inferred from sensory input, and where only some states of the world are under the control of the agent – active inference takes a particular form. Here, the generative model is specified by writing down plausible or allowable policies, hidden states of the world (that must be inferred from observations), and observable outcomes (i.e., sensory input or lower level representations), as well as several matrices that define the probabilistic relationships between these quantities. This sort of generative model is illustrated in the top left panel of figure 1, that describes the model used in this work.

**Figure 1.**
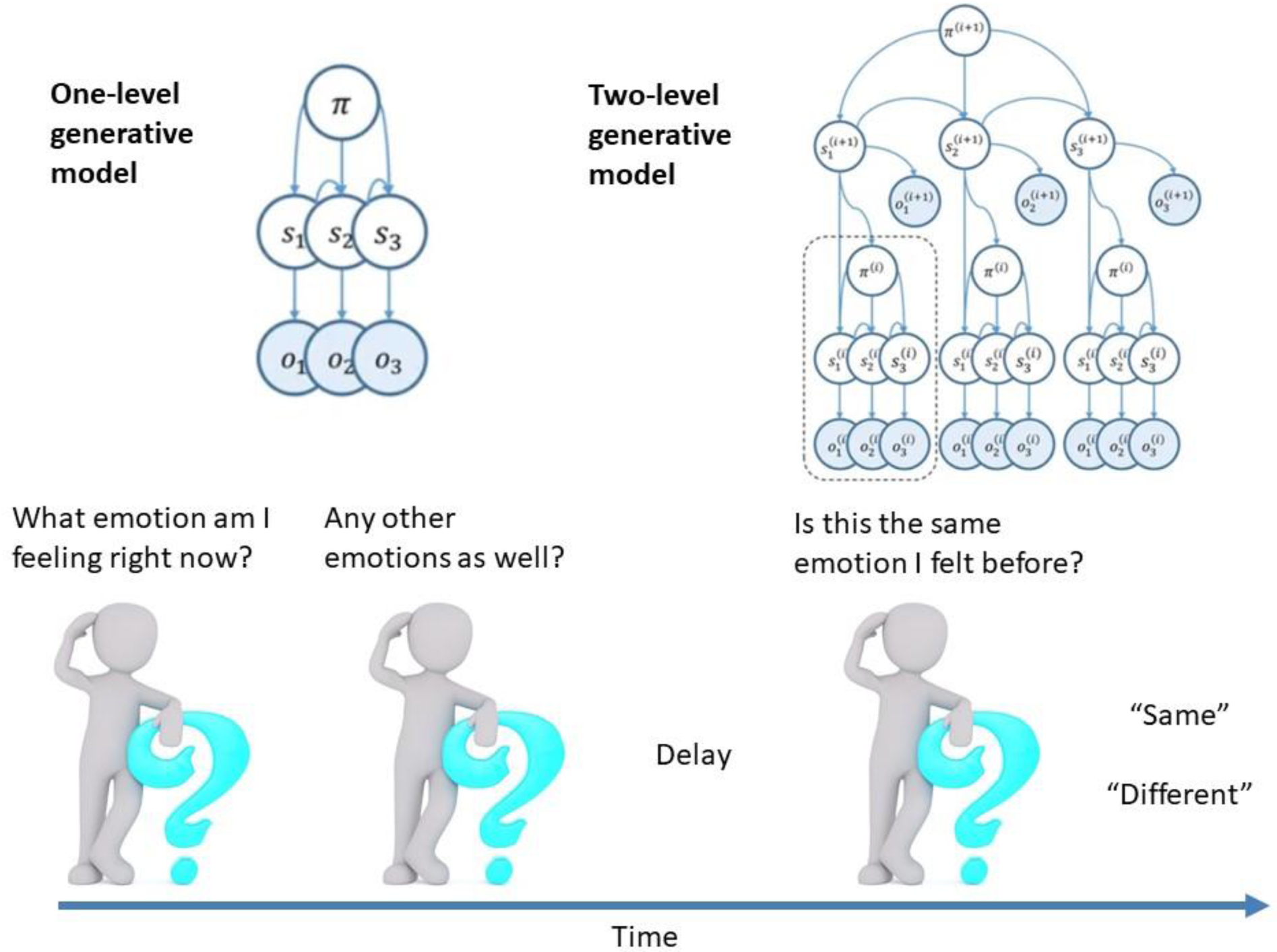
Bottom: Illustration of the working memory task performed by the agent. On each trial, the agent was required to identify the internal state (emotional and/or somatic state) that she is currently feeling at three different points in time. The third internal state that the agent was asked to identify came after a delay period, and the task was to reflect on whether this third internal state was the same as one of the previous two states. Each possible internal state was probabilistically associated with a unique combination of internally observable bodily (valence, arousal, motivation) and cognitive (beliefs about context) features, and the agent needed to selectively attend to these features in order to infer her internal state. Thus, there were in fact two nested tasks – a lower level state recognition task and a higher level working memory task that depended on state recognition. Top left: Illustration of the Markov decision process formulation of active inference used in the simulations described in the main text. The generative model is here depicted graphically, such that arrows indicate dependencies between variables. Here observations (*o*) depend on hidden states (*s*), as specified by the **A** matrix, and those states depend on both previous states (as specified by the **B** matrix, or the initial states specified by the **D** matrix) and the sequences of actions/policies (*π*) selected by the agent. The probability of selecting a particular policy in turn depends on the expected free energy (**G**) of each policy with respect to the prior preferences (**C** matrix) of the agent. The degree to which expected free energy influences policy selection is also modulated by a prior policy precision parameter (**γ**), which in turn depends on beta (***β***) – where higher values of beta entail lower confidence in policy selection (i.e., less influence of the differences in expected free energy over policies). This model, in the single-level form depicted, was used to model the internal state recognition process within the task. Top right: illustration of the two-level generative model used to model the higher level working memory processes: here, the internal states recognized at the first level of the model are treated as the observations made at the second level of the model. This entails a temporally deep structure in which the second level of the model operates at a slower time scale than the first. In this case, the higher level observes and integrates information about the different internal states that are inferred by the lower level at several distinct time points. For more details regarding the associated belief updating, see figure 2 as well as (KJ Friston, Lin, et al., 2017; KJ Friston, Parr, et al., 2017).

The **A**-matrix indicates which observable outcomes are generated by each combination of hidden states (e.g., the likelihood mapping specifying the probability that a particular set of sensory inputs would be observed given a particular set of possible causes outside of the brain). The **B**-matrix is a probability transition matrix, indicating the probability that one hidden state will change into another over time. The agent controls some of these transitions (e.g., those that pertain to the positions of its body) based on the selected policy. The **D**-matrix encodes prior expectations about the initial hidden state the agent will occupy. The **E-**matrix encodes prior expectations about the policies an agent will select, where sufficiently strong expectations for a given policy mean that that policy will be selected habitually (i.e., in a manner insensitive to explicit predictions about future outcomes). Finally, the **C**-matrix specifies prior preferences over observations; it quantifies the degree to which different observed outcomes are preferred or aversive to the agent. In these models, observations and hidden states can be factorized into multiple outcome *modalities* and hidden state *factors*. This means that the likelihood mapping (the **A**-matrix) can also model the interactions among different hidden states when generating outcomes (observations).

In a hierarchical setting where a model has multiple levels (see figure 1, top right panel), the hidden states inferred at the first level are then treated as the observed outcomes at the 2^nd^ level (Friston et al., 2018). These models can also have a deep temporal structure, such that the hidden states at the 2^nd^ level can generate sequences of hidden states at the first level. In the case of reading, for example, the first level of a model could be used to infer the identity of a particular letter, whereas the 2^nd^ level could infer the identity of a word based on a sequence of the letters inferred at the level below.

One potential empirical advantage of active inference model stems from the fact that they have a well-articulated plausible biological basis that affords testable neurobiological predictions. Specifically, these models have companion micro-anatomical neural process theories, based on commonly used message-passing algorithms (Friston et al., 2017a; Parr et al., 2019; Parr and Friston, 2018). Under these process theories, the activation level of different neural populations (typically portrayed as consisting of different cortical columns) encode posterior probability estimates over different hidden states (see figure 2). These activation levels are then updated by synaptic inputs with particular weights that convey the conditional probabilities encoded in the **A** and **B** (among other) matrices described above. In what follows, we describe how a hierarchical generative model was specified to produce an agent with different levels of emotional awareness. We then provide examples of the types of simulated neuronal responses that can be generated – and potentially lead to empirical predictions that could be tested in future clinical neuroscience research.

**Figure 2.**
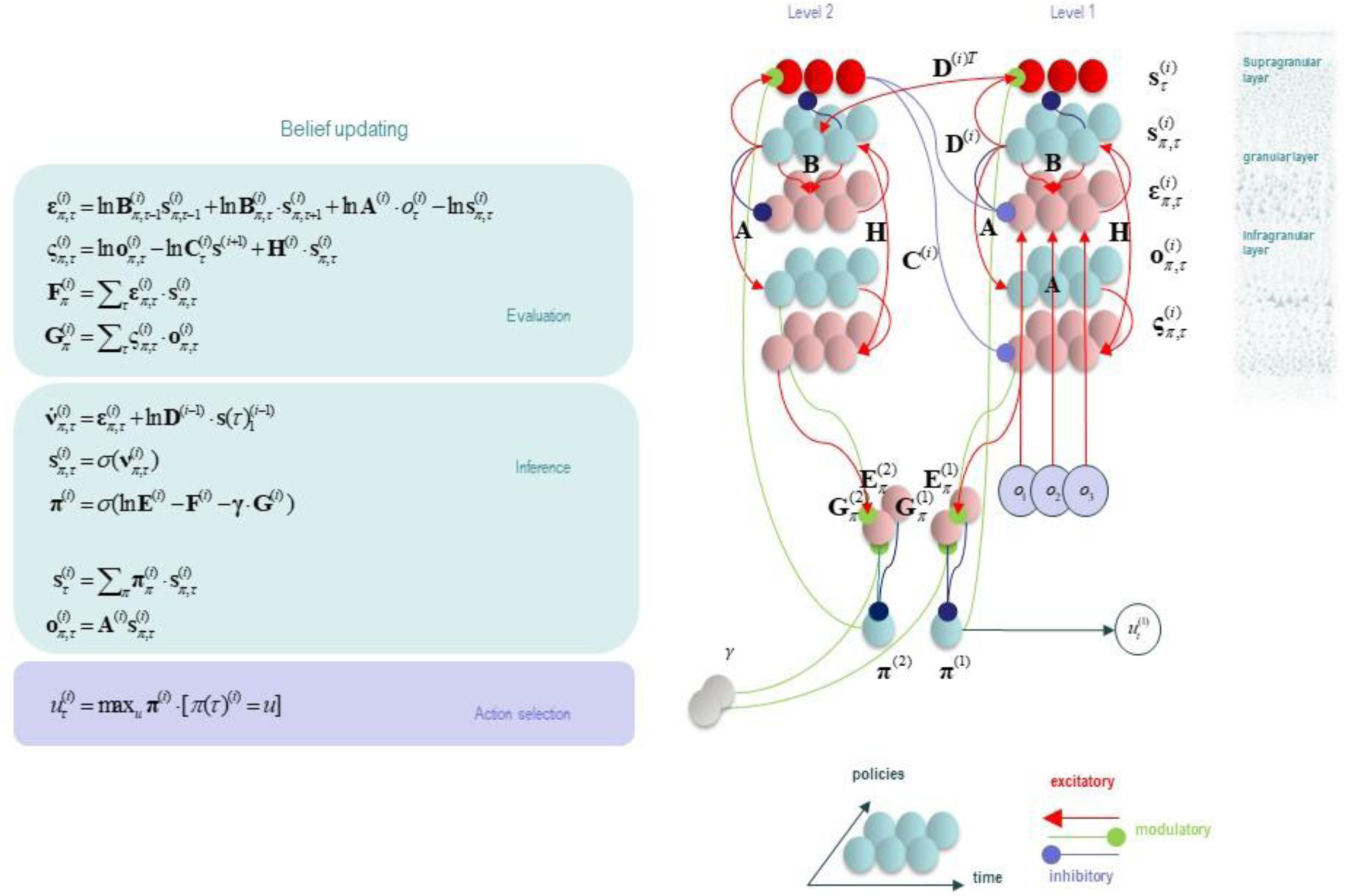
This figure illustrates the neuronal message passing framework for the hierarchical MDP formulation of active inference that was used to perform the simulations described in this paper. The differential equations in the left panel approximate Bayesian belief updating within the graphical model depicted in the upper portion of Figure 1 via a gradient descent on free energy (**F**). The right panel illustrates the proposed neural basis by which neurons making up cortical columns at two interconnected hierarchical levels could implement these equations. The equations have been expressed in terms of two types of prediction errors. State prediction errors (**ε**; line 1) signal the difference between the (logarithms of) expected states (**s**) under each policy and time point—and the corresponding predictions based upon outcomes/observations (**A** matrix) and the (preceding and subsequent) hidden states (**B** matrix, and **D** matrix for the initial hidden states at the first time point). These represent prior and likelihood terms respectively – depicted as messages being passed between neural populations (colored balls) via particular synaptic connections in the right panel. These (prediction error) signals drive depolarization (**v**; line 5) in the neurons encoding hidden states (**s**), where the probability distribution over hidden states is then obtained via a softmax (normalized exponential) function (**σ**; line 6). Outcome prediction errors (**ς**; line 2) instead signal the difference between the (logarithms of) expected observations (**o**) and those predicted under prior preferences (**C**). This term additionally considers the expected ambiguity or conditional entropy (**H**) between states and outcomes (computed from the **A** matrix). This prediction error is weighted by expected observations (line 9) to evaluate the expected free energy (**G**; line 4) for each policy (**π**). These policy-specific free energies are then integrated to give the policy expectations via a softmax function (line 7). The **E** matrix in line 7 reflects a prior distribution over policies (which can be thought of as encoding habits based on self-observed regularities in past behavior), and γ is a policy precision parameter that modulates the degree to which policy selection is influenced by habits (**E**) vs. model-based beliefs (**G)**. Actions at each time point (**u**; line 10) are then chosen out of the possible actions under each policy weighted by the value (negative expected free energy) of each policy. Line 8 expresses a Bayesian model average that is necessary for estimating the probability over states (by taking into account the probability of each policy and the probability of each state given each policy). The resulting posterior distributions over states at the first level are then treated as observations by the second level (levels are denoted by the superscript *i*, where each level operates based on the same equations but over different timescales). In the proposed neuronal implementation on the right, probability estimates have been associated with neuronal populations that are arranged to reproduce known intrinsic (within cortical area) connections. Red connections are excitatory, blue connections are inhibitory, and green connections are modulatory (i.e., involve a multiplication or weighting). These connections mediate the message passing associated with the equations on the left panel. Cyan units correspond to expectations about hidden states and (future) outcomes under each policy, while red states indicate their Bayesian model averages. Pink units correspond to (state and outcome) prediction errors that are averaged to evaluate expected free energy and subsequent policy expectations (in the lower part of the network). Expected free energy and policy evaluation are each depicted as occurring subcortically. This (neural) network formulation of belief updating means that connection strengths correspond to the parameters of the generative model described in the text. Only exemplar connections are shown to avoid visual clutter. Furthermore, we have just shown neuronal populations encoding hidden states under two policies over three time points (i.e., two transitions), whereas in the task described in this paper there are greater number of allowable policies. For more information regarding the mathematics and processes illustrated in this figure, see (KJ Friston, Lin, et al., 2017; KJ Friston, Parr, et al., 2017).

## 3. A deep temporal model of emotional awareness

In this paper, we explicitly model the second and third processes in the TPM: affective response representation (emotion conceptualization) and conscious access. Within these processes, after being generated in a particular context, the (exteroceptive, proprioceptive and interoceptive) aspects of an affective response are first used to infer the current emotional state, and representations of this emotional state are then made available to domain-general cognition and held in working memory, where they can be combined with other information and used to inform decision-making and verbal reporting. The emotion conceptualization process is necessary for the third level of emotional awareness in the theory of levels of emotional awareness (Lane et al., 1990), where an individual is capable of becoming aware of single emotional states like sadness and fear. Note that an emotional state is a hidden state in the generative model, meaning that emotions play the role of hypotheses or representations that best explain the multimodal sensations at hand (e.g., inferences corresponding to contents such as “I am anxious” are implicitly encoded as the best explanations for particular patterns of current exteroceptive and interoceptive signals). The conscious access process is necessary for the fourth and fifth levels of emotional awareness in the theory, where an individual is capable of holding information about emotions in mind over an extended period of time, combining it with other information, and using this integrated information within domain-general cognition – as in the ability to contemplate multiple emotions at once or the ability to simultaneously think about one’s own emotions and someone else’s (e.g., “She is anxious because I am clearly frightened”).

To model this type of emotion-focused cognition, we constructed a simple working memory task for an agent or synthetic subject to perform (see figure 1, bottom panel). In this task, the agent was presented with different affective bodily responses – characterized by distinct combinations of valence, arousal, and action tendencies – associated with particular (interpretations of) contexts. After the onset of each affective response, the agent’s task was to attend to the different aspects of this response and its eliciting context, and to categorize that response under different possible internal state concepts (some of which corresponded to emotion concepts and others which corresponded to non-emotional somatic concepts). Subsequently, however, the agent was also instructed to hold that internal state in working memory, while experiencing a second affective response. This was followed by a delay period, after which the agent experienced and categorized yet a third affective response. While holding all three perceived internal states in mind, the agent’s task was to self-report whether the third internal state was the same or different in relation to one of the first two states. This particular task was chosen for two reasons: 1) because of its similarity to previously published studies of emotion-focused working memory and its relation to emotional awareness (R Smith et al., 2017b; Smith et al., 2018b, 2018c; Waugh et al., 2014; Xin and Lei, 2015), and 2) because it requires several elements associated with different levels of emotional awareness; including perceived bodily responses, internal state categorization, and the ability to hold multiple internal states in mind at once.

To simulate affective responses, internal state categorization, and working memory (based on the task described above), we first needed to specify an appropriate generative model. Once this model is specified, standard (variational) message passing schemes can be used to simulate belief updating and behavior in a biologically plausible manner (for more detail, please see figure 2 as well as (Friston et al., 2017a, 2017b)). In the first level of our model, the observable aspects of an affective response (the outcomes in the model) included a feature space including: two possible valences (neutral, unpleasant), two possible interoceptive arousal levels (low, high), two possible motivational states (approach, avoid), and three types of beliefs about the eliciting context (i.e., a belief that it is a non-threatening context, social threat context, or physical threat context).

It is worth highlighting that these features are fairly high-level observations, which would need to be derived from interoceptive and exteroceptive sensory processing at lower levels; i.e., where these sensations would themselves have been generated by a stimulus/context and the subsequent affective response generation process of the TPM, which we do not explicitly model here. To model the belief updating of interest here, these affective response features were sufficient for our purposes. Thus, in the simulations below, we simply presented the model with affective and contextual cues to assess the degree of awareness attained by the agent. We do not model the generation of affective bodily outcomes themselves (i.e., which would occur via selection of visceromotor and skeletomotor policies) in response to a context and its subjective interpretation. In short, we assume affective outcomes reflect inferences drawn by a lower hierarchical level, whose role is to explain interoceptive data: e.g., as in (Allen et al., 2019). This means that they should not be interpreted as sensory data, but as lower-level inferences about the causes of sensory data. For example, we might expect changes in valence to correspond to changes in the precision/confidence associated with lower-level visceromotor and skeletomotor policy selection, or to related internal estimates that can act as indicators of a model’s current successfulness in resolving uncertainty and achieving preferred outcomes; see (Clark et al., 2018; de Berker et al., 2016; Joffily and Coricelli, 2013; Peters et al., 2017)).

In the first level of our model (see figure 3A and 3B), the **A**-matrix specified a likelihood mapping between affective outcomes and the first hidden state factor, which included five internal state concepts that could explain outcomes (neutral state, sadness, panic, sickness, heart attack), such that each outcome combination was predicted by each concept category with distinct probabilities. Note, we focus primarily on negatively valenced states here, due to their clinical relevance and relation to somatic misattribution. These mappings meant that: 1) an emotionally neutral state predicted neutral valence, either low or high arousal, either approach or avoidance motivation, and a non-threatening context; 2) sadness predicted negative valence, low arousal, avoidance, and a social threat context; 3) panic predicted negative valence, high arousal, avoidance, and a social threat context; 4) sickness predicted negative valence, low arousal, avoidance, and a physical threat context; 5) heart attack predicted negative valence, high arousal, avoidance, and a physical threat context.

**Figure 3.**
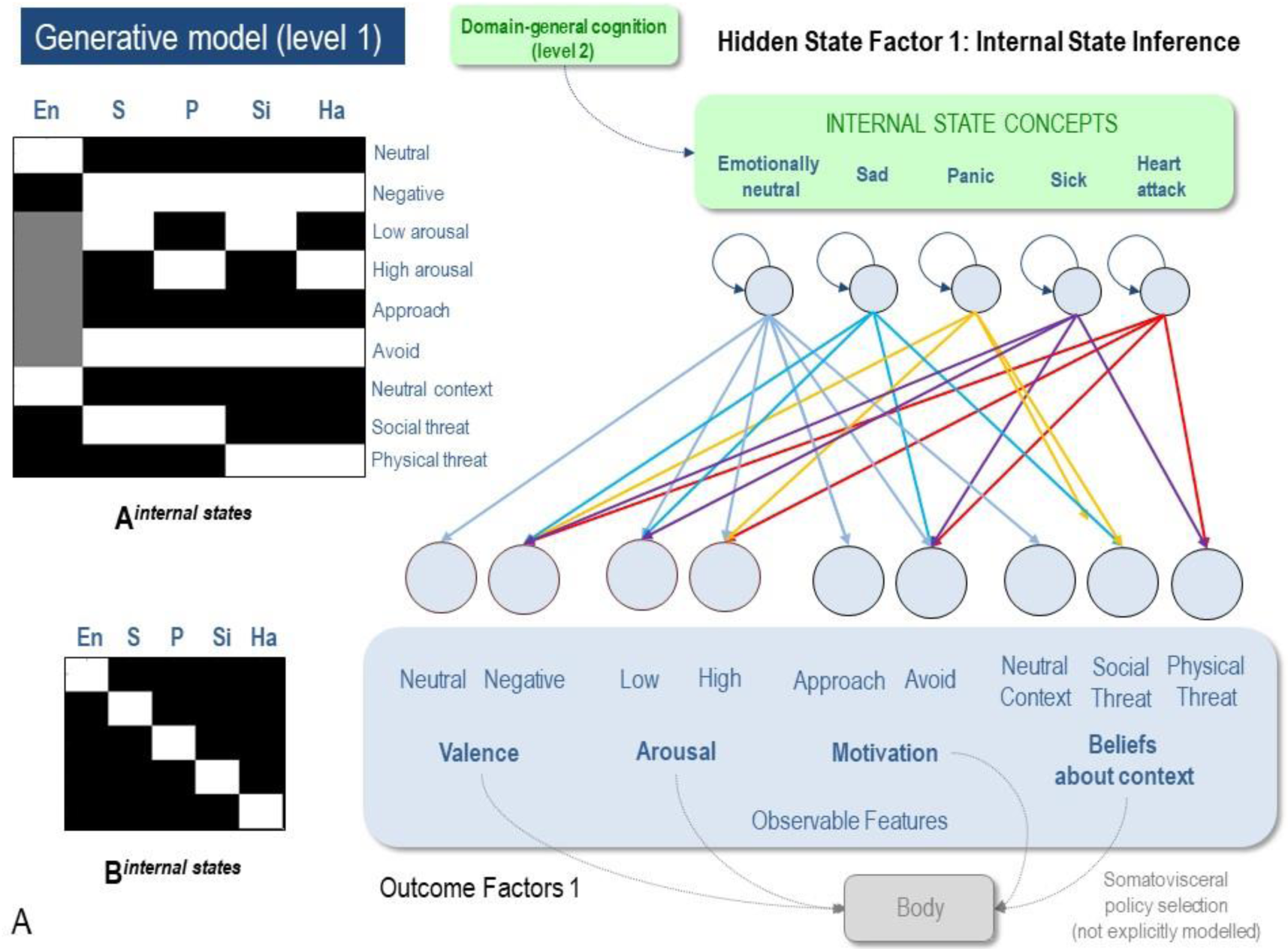
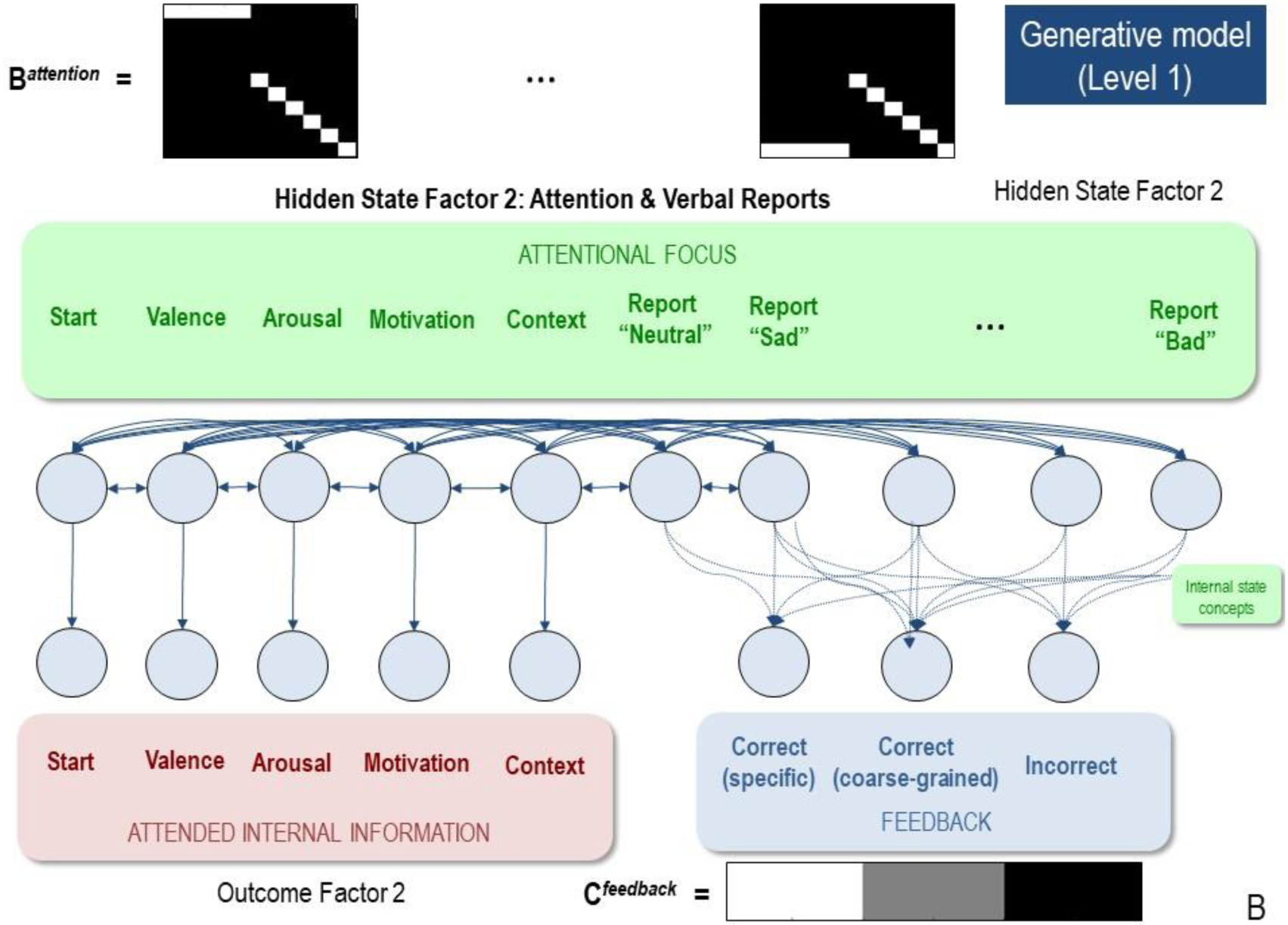
First-level model. (A) Displays the levels of hidden state factor 1 (internal state concepts) and their mapping to different lower-level representations (here modeled as affective outcomes and feedback). Each internal state generated different outcome patterns with different probabilities (see text for details). The **A**-matrix encoding these mappings is illustrated in the upper left (i.e., lighter colors indicate higher probabilities). The **B**-matrix in the lower left illustrates an identity mapping between internal states, such that internal states were believed, *a priori*, to be stable within trials. The precision of these matrices (i.e., implicit beliefs about how precisely different internal states are associated with different observations or beliefs about internal state stability) could be adjusted via passing them through a softmax function with different precisions. This model simulates the affective response representation process within the three-process model of emotion episodes (Smith et al., 2018a). The grey arrows/boxes at the bottom of the figure denote a further process within the three-process model (i.e., affective response generation, understood as a form of lower-level skeleto/visceromotor policy selection) that is not explicitly modeled in the current work. (B) Displays the levels of hidden state factor 2 (focus of attention) and its mapping to outcomes. Each focus of attention mapped deterministically (the **A**-matrix was a fully precise identity matrix) to a “location” (i.e., an internal source of information) at which different outcomes could be observed. The **B**-matrix in the upper left illustrates that the agent could choose to shift her attention from one internal representation to another to facilitate inference. The final attentional shift in the trial was toward a (proprioceptive) motor response to report an internal state (i.e., at whatever point in the trial the agent became sufficiently confident, at which point the state could not change until the end of the trial), which was either correct or incorrect. The agent most preferred (expected) to be correct in reporting a specific internal state and least preferred to be incorrect. If the agent was not sufficiently confident to report a specific emotion, but was confident that it was feeling some type of negative emotion, it could also choose to simply report the more coarse-grained feeling of “bad” – which was preferred to an intermediate degree between the other two possible outcomes. Because policies (i.e., sequences of implicit attentional shifts and subsequent explicit reports) were selected to minimize expected free energy, inference under this model induces a sampling of salient representational outcomes and subsequent report – under the prior preferences described above.

While certainly oversimplified, the likelihood mappings specifying the content of sadness and panic were motivated by previous literature linking these emotion concepts to the above-mentioned lower-level outcomes. For example, the concept of sadness, as modeled here, is based on research (e.g., (Badcock et al., 2017)) focused on the low-arousal form of sadness that is more often associated with social isolation and clinical depression (i.e., as opposed to a higher-arousal, acute form of sadness associated with active crying, etc.). The concept of panic, as modeled here, is based primarily on the types of panic attacks that often occur in social contexts (e.g., crowded public spaces) and that are often associated with agoraphobia and/or social anxiety disorder (Jack et al., 1999; Potter et al., 2014). Of course, panic attacks can also occur in objectively dangerous situations, but more often there is no objective physical danger and helping an individual to identify the affective origin of panic-related bodily sensations can itself aid in emotion regulation (e.g., “okay, this is just a panic attack – it will go away in a few minutes and I’m not going to die”).

It is also important to clarify that these likelihood mappings are probabilistic; for example, one could still feel neutral at a funeral or feel sick at a crowded event. These possibilities were simply coded as having lower probabilities. The specific precision of these probabilistic mappings was controlled via the precision or inverse temperature parameter of a softmax function applied to a fully precise form of the likelihood mappings between concepts and affective outcomes; for a more technical account of this type of precision or attention modelling, please see (Parr and Friston, 2017a)). By default, the precision applied to these mappings (and to the working memory content mappings at the higher level described below) was set to a value of 1.5, resulting in a relatively precise mapping that was sufficient for high performance (however, lower precisions were applied to specific state-outcome mappings in specific simulations described below). The corresponding **B-**matrix for the internal state factor was an identity matrix, such that the internal state was *a priori* stable across each trial, with greater or lesser precision.

To incorporate the role of selective attention in emotional awareness – a type of mental action (Limanowski and Friston, 2018; Metzinger, 2017) – we also included a state corresponding to *attentional focus*. Specifically, precise information about the various aspects of an affective outcome was only available within certain attentional states (e.g., the agent needed to pay attention to her bodily arousal to learn whether it was high or low). This attention-dependent access to information was built into the **A-**matrix mapping internal states to different affective outcomes (e.g., arousal level) under different attentional states, such that these mappings were only informative when the agent adopted the associated attentional state (e.g., the “attention to arousal” state). The **B-**matrices for this factor specified all possible (controllable) transitions between attentional states, such that the agent could choose all one-step policies that allowed her to attend to as few or as many features as she deemed necessary, prior to reporting her beliefs about her own internal state. Self-reports were also modeled as additional states within the same hidden state factor, such that the agent can attend to emotional states *or* make a report once sufficiently confident (but not both). This included reporting the belief that she was feeling neutral, feeling sad, feeling panic, feeling sick, or experiencing a heart attack. If insufficiently confident about which emotion was experienced, she could also simply report feeling emotionally bad. At this point, the agent would also observe a type of “social feedback” indicating whether the self-reported state matched the hidden state that generated the observed pattern of affective response features. The **C**-matrix was constructed such that the agent preferred to correctly report one of the specific internal states, least preferred to receive feedback that she was incorrect, and had an intermediate preference for being correct about the more general state of feeling bad. To ensure that the agent was sufficiently motivated to attend to multiple features before selecting a report, the agent assigned higher negative value to reporting incorrectly than she assigned positive value to observing correct feedback.

The second level of our model corresponded to domain-general cognition (see figure 4A and 4B). This level included 4 hidden state factors. The first was the state to be remembered where we included 6 exemplary states: sadness, panic, both sadness and panic, both sadness and sickness, both heart attack and panic, and both heart attack and sickness. The reason that some of these are composite, while others are single states, relates to the trial structure outlined below. In short, there are two emotional state presentations before the delay. These may include two different states or a neutral state and a single emotional state. While we could have included all possible combinations of first level states, these were sufficient to simulate the relevant mechanisms associated with emotional awareness described below. They can also be mapped onto clinically relevant cognitive states; for example, I could recognize that I am sad because I am sick, or that I am sad because I just had a terrible panic attack in public. The second hidden state factor corresponded to the 3^rd^ internal state presented to the second hierarchical level, to which she compared the first two (and answered whether it was the same state or a different state from the first two). This can be thought of as modelling a situation in which I reflect upon whether my current emotional state is similar to how I felt at a particular point in the past (e.g., as often occurs in psychotherapy contexts). The third hidden state factor was the time point within the trial. This included the starting state, the first internal state, the second internal state, a maintenance phase, the third internal state, and then the response phase. The fourth hidden state factor was the agent’s report (undecided, same, different).

**Figure 4.**
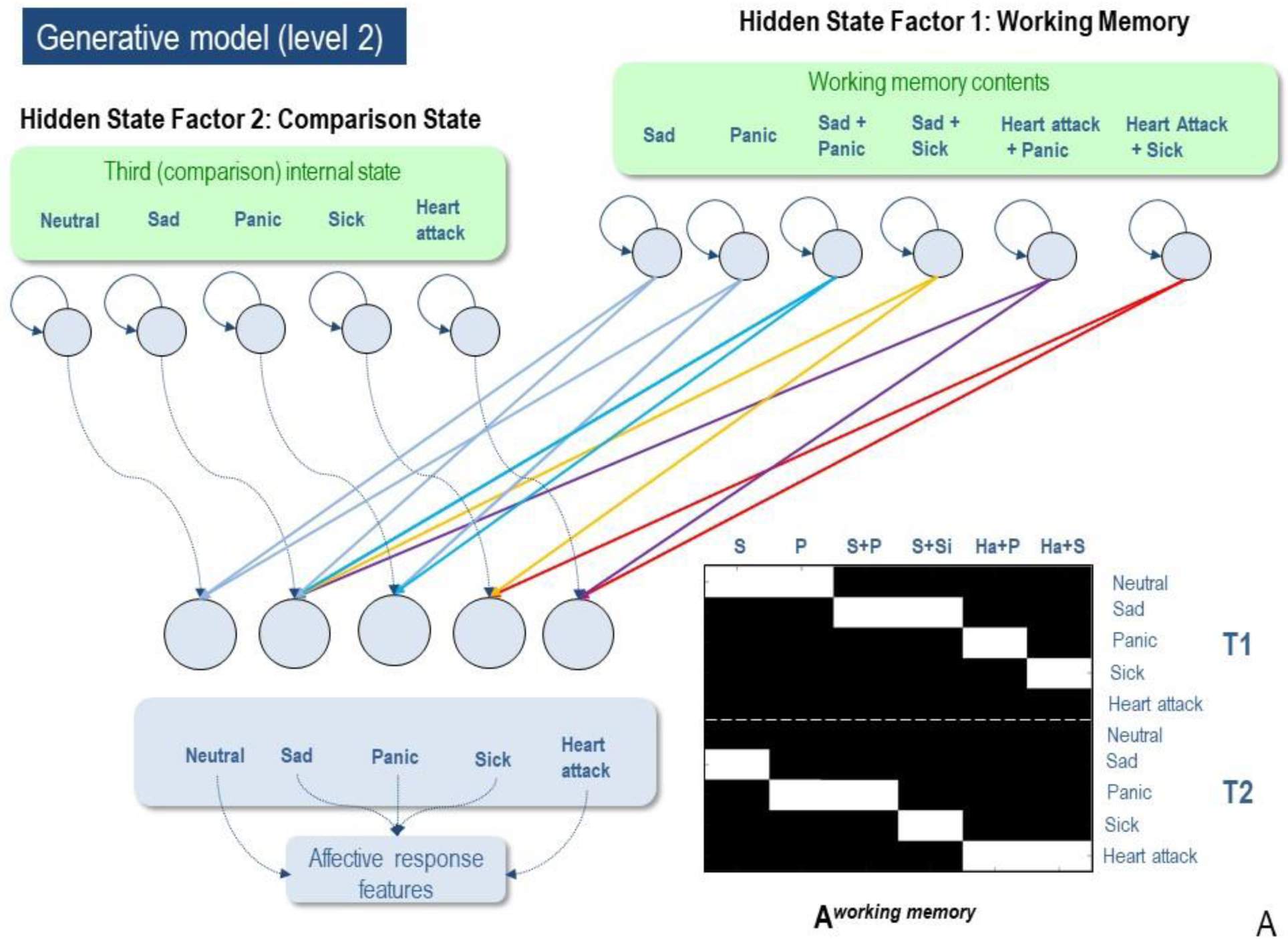
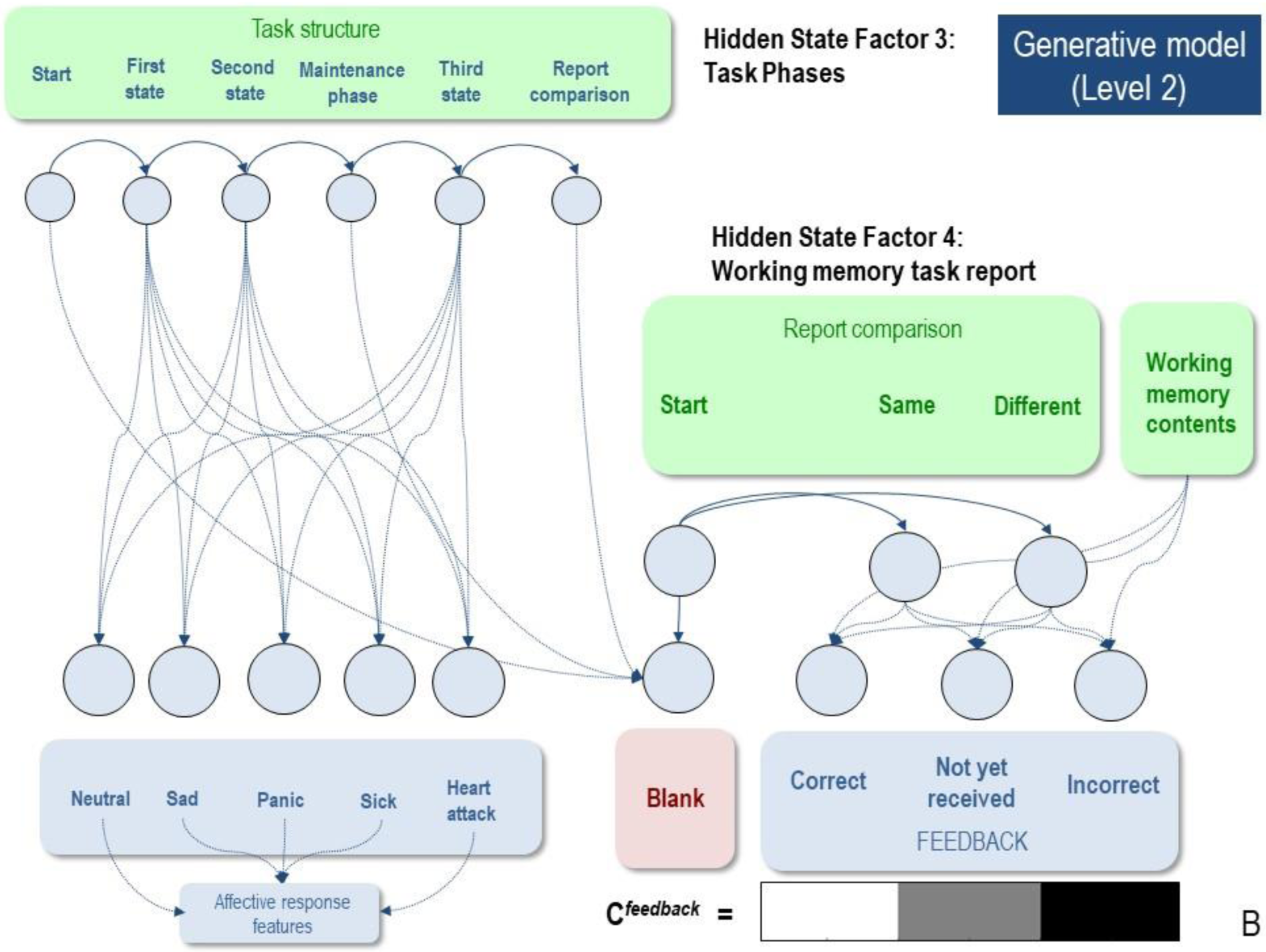
Second-level model. (A) Displays the levels of hidden state factor 1 (working memory contents) and 2 (the third comparison state), and their mapping to the different internal states at the lower level. Each possible emotional context corresponded to different combinations of internal state representations (see text for details). The colored arrows illustrate these mappings as they are specified in the **A**-matrix, which is also depicted in the bottom right. The **B**-matrix specified an identity mapping between working memory contents, such that contents were stable within trials. The precision of these matrices (i.e., implicit beliefs about how precisely different internal state representations update working memory contents or about the stability of working memory contents over time, respectively) could be adjusted via passing them through a softmax function with different precisions. This higher-level model simulates the conscious access process within the three-process model of emotion episodes (Smith et al., 2018a). (B) Displays the levels of hidden state factor 3 (task structure) and 4 (self-report), and their mapping to both internal states and feedback. Each of these mappings were fully deterministic (the relevant **A**-matrices were maximally precise). The relevant **B**-matrices specified deterministic transitions from one task phase to the next and allowed the agent to report that the third internal state was either the “same” or “different” in relation to the previous two internal states experienced (i.e., deep policies in which the agent could only remain in an “undecided” state until the final report phase at which time it could make its choice). The agent most preferred (expected) to be correct.

The first outcome modality corresponded to each of the five possible internal states that were inferred at the lower level, as well as a “blank” outcome that corresponded to the starting state and the maintenance phase (this blank outcome was included as an additional hidden state in the lower-level model described above, but had imprecise mappings to affective outcomes; this is omitted in our figures depicting the first-level model for clarity). The second outcome modality corresponded to feedback, such that after each trial the agent was told whether she was correct or incorrect in the comparison between the three internal states experienced over time. The **C**-matrix was then set such that the agent preferred to observe correct feedback, and available policies only included choosing to report either “same” or “different.”

## 4. Simulating individual differences in emotional awareness

### 4.1 Foundational simulations

As an initial validation, we enabled the first level of the model and presented it with 100 different internal state responses – 20 corresponding to each of the five internal state concepts. With a moderately precise mapping from internal state concepts to affective outcomes (precision = 1.5), we observed that the agent correctly inferred and reported her internal state 100% of the time. This confirmed that, with moderately precise probabilistic state-outcome mappings, the agent could successfully deploy selective attention and infer internal states (see figure 5A for illustration of an example trial).

**Figure 5.**
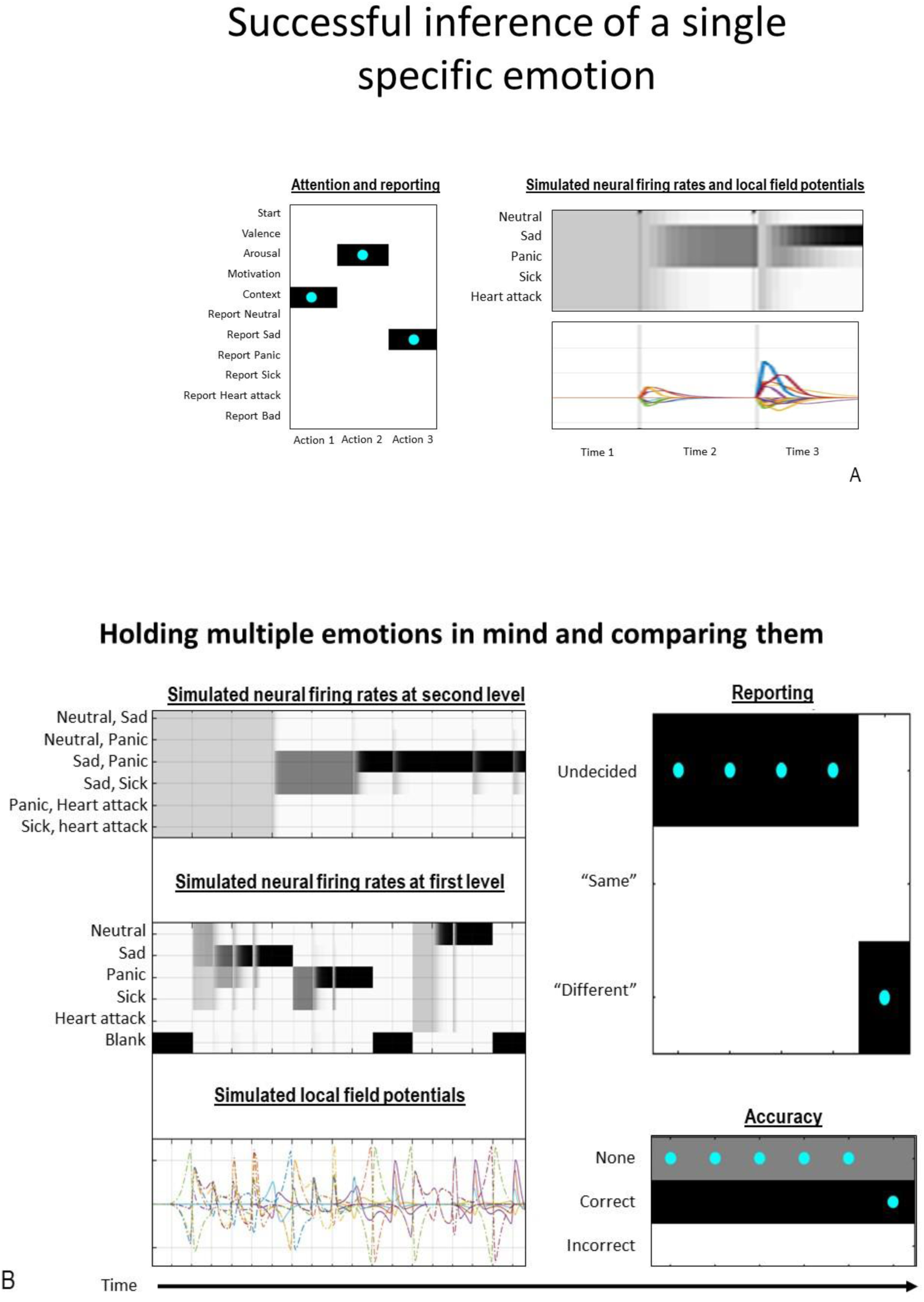
Example trials illustrating optimal performance under high levels of emotional awareness. (A) shows successful inference about a single emotional state in the lower level of the model. As shown on the left, the agent first chose to attend to her beliefs about context (and observed social threat) and then chose to attend to her arousal level (and observed low arousal), at which point she was sufficiently confident to report feeling sad (cyan dots indicate the true action taken; darker colors indicate higher levels of confidence in one action over others). The right panel illustrates the simulated neuronal firing rates (darker = higher firing rate) and local field potentials (rates of change in firing rates) that would be predicted under the neural process theory associated with active inference that is depicted in figure 2 (Friston et al., 2017a). (B) Displays successful inference of a combination of two emotions in the higher-level model. The right panel shows that the agent reported “different” and then observed “correct” feedback. The left panel shows simulated neuronal firing rates and local field potentials for both the first and second levels of the model. As can be seen, at the first level, the first emotion represented was sadness and the second emotion represented was panic, followed by a delay (“blank” state), followed by the third represented emotion, which was neutral. At the second level, it can be seen that evidence first accumulates to suggest that sadness and panic are both present, at which point firing rates in the neural populations encoding that combination are maintained throughout the rest of the trial so that they can be used to inform decision-making at the end.

We then engaged the second level of our model, and presented it with multiple internal states over time, corresponding to each of the six internal state combinations included in the model. Here, we presented the model with 10 examples of each of the six internal state combinations (i.e., 60 trials in total). We again observed that the model was capable of successfully gating each of these combinations of internal states into working memory, holding them in an active state over a delay period, and reporting correctly whether a third internal state matched one of the previous two (see figure 5B for an example trial of holding both sadness and panic in mind). Thus, the model performed optimally, both at inferring its own internal states and subsequently holding them in memory to perform subsequent cognitive operations on them. The model in this particular configuration was therefore capable of emulating emotional awareness.

Having constructed our model and establishing that it could generate emotional awareness behavior, we will now consider a number of exemplar simulations demonstrating that distinct mechanisms can produce different trait levels of emotional awareness – as it is empirically measured via reporting behavior. As discussed in the introduction, articulating these distinct mechanisms may be clinically useful, as they imply distinct therapeutic intervention targets. They also highlight distinct potential mechanisms that could be used as hypotheses to guide the development of measures that could phenotype individuals in terms of the underlying processes contributing to low levels of emotional awareness.

To illustrate different types of aberrant belief updating that would produce somatic misattribution (EA level 1) and low emotion granularity (EA level 2), as well as belief updating allowing for awareness of single granular emotions (EA level 3), we will focus on the first level of our model. Note that, for illustrative purposes, in these simulations we allow the agent to report her emotions in the absence of first gating them into domain-general cognition at the second level. However, as there is behavioral evidence that emotion concepts can be primed in the absence of awareness (Smith and Lane, 2016; Zemack-Rugar et al., 2007), and a large body of neural and behavioral evidence that the brain can represent information in the absence of conscious awareness (Dehaene, 2014), actual self-reportability is thought to further depend on the higher-level conscious access processes included in the second hierarchical level of our model. Thus, we will subsequently focus on the second (working memory) level of our model, which is relevant to understanding individual differences in the ability to hold single or multiple emotions in mind over an extended period of time (i.e., a necessary condition for EA levels 4 and 5).

### 4.2 Mechanism 1: Abnormal affective response generation

Although fairly simple, one straightforward mechanism that would lead to the absence of self-reported emotions is if affective outcomes are not available in the first place. Somewhat trivially, if our model is only presented with neutral outcomes, it never reports awareness of any emotions – even if it has precise emotion concept knowledge. Although seemingly trivial, this potential mechanism is worth highlighting, because it appears to be prescient for some individuals with low emotional awareness (and high alexithymia scores; (Smith et al., 2019b)). Such individuals can show an absence of normative skin conductance responses as well as atypical valence-related facial muscle responses when presented with affective stimuli. Yet, they can understand emotion concepts and perform well at recognizing the emotions of others.

The cause of this type of deficient affective response is unclear. One possibility is that it could reflect congenital abnormalities in the neural circuitry associated with the generation of unconditioned and/or conditioned bodily responses to normatively emotion-provoking stimuli. Another possibility, however, is that it relates to the way in which perceived situations are cognitively evaluated in terms of their significance to the individual. Within appraisal theories of emotion, for example, affective responses are generated based on appraisal dimensions such as the controllability of a situation, whether or not it is congruent with one’s goals and values, whether it was expected or unexpected, whether one assigns responsibility to the self or others, among others (Moors et al., 2013; Scherer, 2009). Thus, affective responses could also fail to be generated if such appraisal processes and related mechanisms for predicting present and future metabolic demands failed to process information in a normative and adaptive manner.

### 4.3 Mechanism 2: Strong prior expectations for somatic conditions

The second mechanism in our model that could plausibly produce low emotional awareness involves having learned strong prior expectations for dangerous somatic conditions (e.g., as in people who have high levels of anxiety sensitivity or are otherwise preoccupied with possible threats to health; see (Mueller and Alpers, 2006) for evidence that anxiety sensitivity is associated reduced awareness of emotion). To illustrate this, we equipped the model with biased (precise) expectations that the dangerous somatic states of heart attack and sickness were more likely than emotional states. As in our initial model simulations described above, we presented the model with 20 responses corresponding to each of the five internal states – but did so under various levels of somatic expectation. These simulations showed that if model parameters were specified as if the agent expected to experience dangerous somatic states at least seven times more often than emotional states, then affective responses began to be reliably misrecognized as indicative of somatic conditions. Specifically, the agent showed a strong tendency to somatize, misrecognizing sadness as sickness and panic as a heart attack (see figure 6 for an example trial and simulated neuronal responses that would be predicted based on the neural process theory depicted in figure 2). This was due in part to the overlap between these pairs of internal states (e.g., sadness and sickness both involve avoidance, negative valence, and low arousal; panic and heart attack both involve avoidance, negative valence, and high arousal). To differentiate these pairs, it was necessary to attend selectively to contextual factors. However, as illustrated in figure 6, when equipped with beliefs that somatic explanations are more likely, such factors tended to be ignored. Specifically, the agent tended to first attend to arousal, and then “jump to conclusions” that her observations indicated sickness if arousal was low (in 100% of the simulated trials) and heart attack if arousal was high (in 95% of the simulated trials). This form of belief updating may therefore provide a possible explanation for the type of somatic focus and misattribution associated with the lowest level of emotional awareness.

**Figure 6.**
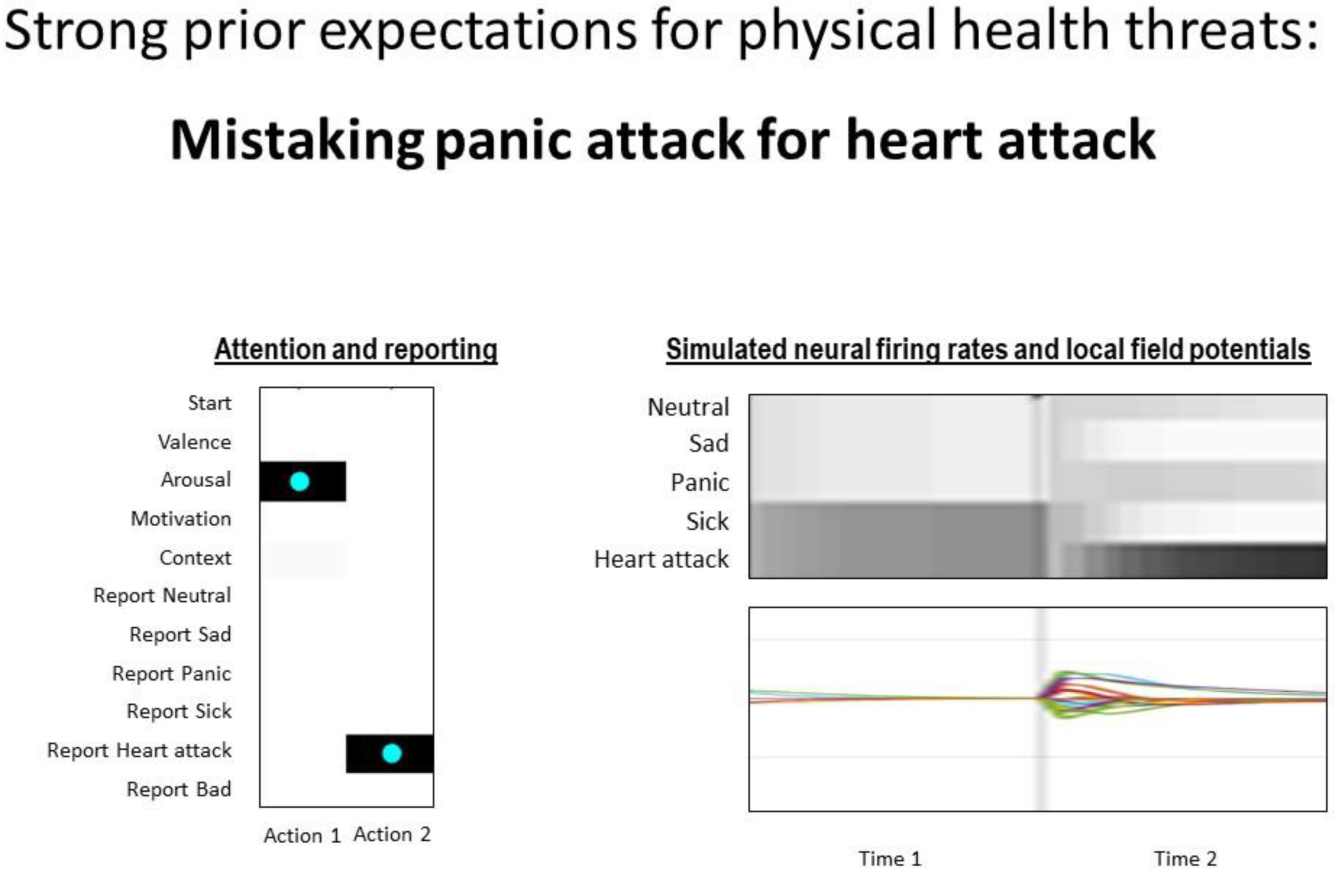
Example trial displaying suboptimal performance as a result of strong prior expectations for the presence of physical health threats (i.e., states such as sickness and heart attack). As shown on the left, the agent first chose to attend to her arousal level (and observed high arousal), at which point it immediately inferred that heart attack was most likely (cyan dots indicate the true action taken; darker colors indicate higher levels of confidence in one action over others). The right panel illustrates the simulated neuronal firing rates (darker = higher firing rate) and local field potentials (rates of change in firing rates) that would be predicted under the neural process theory depicted in figure 2 (Friston et al., 2017a). As can be seen, the neuronal populations encoding sickness and heart attack start out with elevated firing rates, and upon observing high arousal, firing rates further increase in the neuronal population encoding heart attack.

### 4.4 Mechanism 3: Poor conceptual understanding of emotions

A third plausible mechanism that could produce low levels of emotional awareness corresponds to poor emotion concept acquisition – as may occur in individuals who fail to learn about emotions due to impoverished social learning opportunities early in development (e.g., parental neglect; (Lane et al., 2018)). Here, impoverished knowledge about emotions was simulated by reducing the precision of the mapping from emotion concepts to affective response features at the lower level, such that each emotion concept less clearly predicted one pattern of affective response features over others. This was accomplished by passing the **A**-matrix encoding these state-outcome mappings through a softmax function with a low precision (0.01), leading to significantly less precise mappings. Under these conditions, the agent tended to avoid reporting specific emotions, and instead simply chose to report that it felt “bad” in 85% of simulated trials. As exemplified in figure 7, the agent’s uncertainty also promoted repeated shifts in attention in an attempt to gain more information before reporting (leading to slower reactions times). Thus, the primary result of poor emotion concept acquisition was a reduction in granularity, where distinct affective responses were not conceptually differentiated (i.e., level 2 emotional awareness).

**Figure 7.**
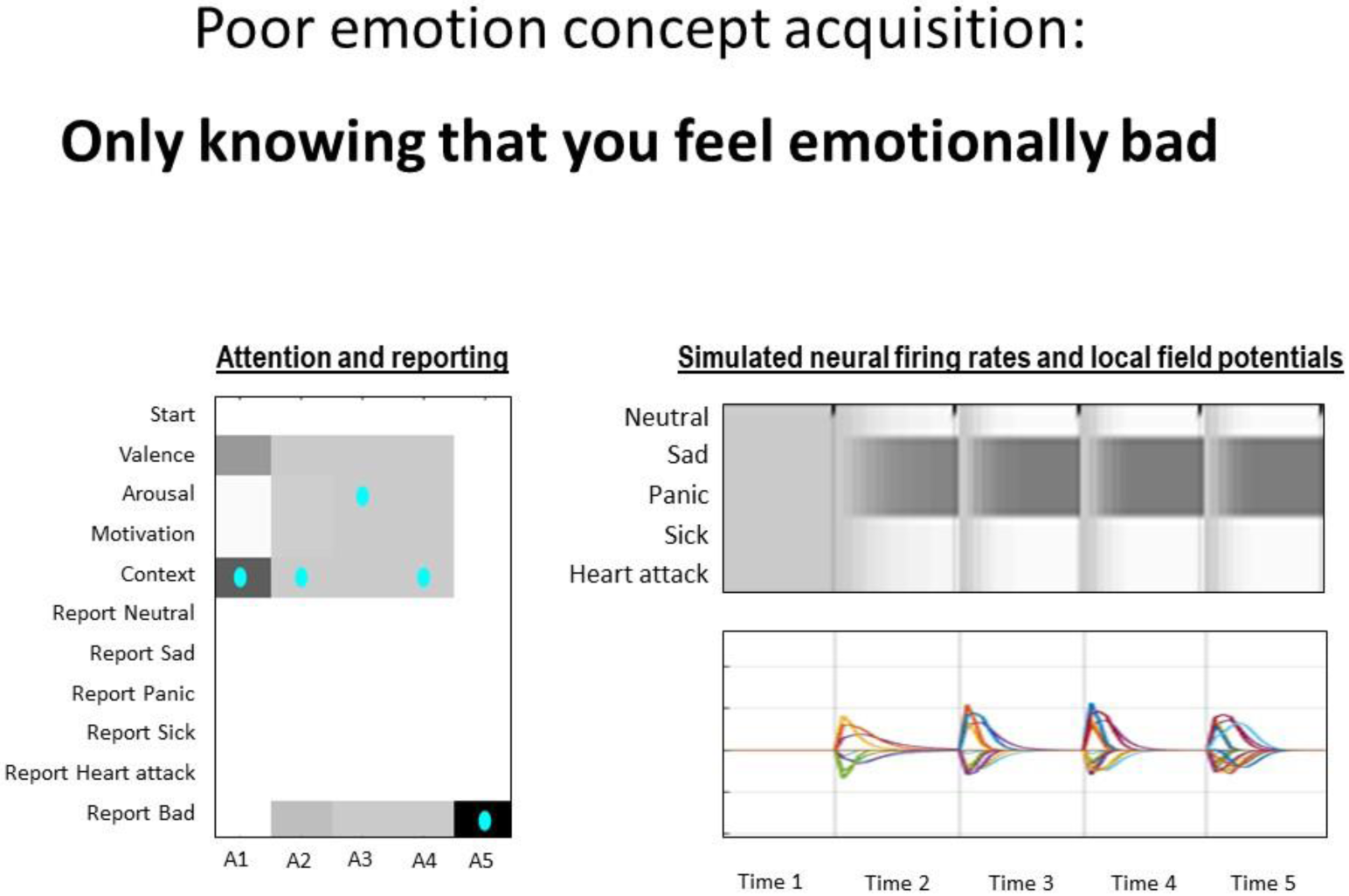
Example trial displaying suboptimal performance as a result of poor emotion concept acquisition (here operationalized as a highly imprecise mapping from emotional state representations to the observable features associated with an affective response at the level below). As shown on the left, the agent first confidently chose to attend to her beliefs about context (and observed social threat), at which point she became highly uncertain about what to attend to next to further reduce her uncertainty and continually shifted between attending to arousal and context information until the final time step, where she simply reported feeling bad (cyan dots indicate the true action taken; darker colors indicate higher levels of confidence in one action over others). The right panel illustrates the simulated neuronal firing rates (darker = higher firing rate) and local field potentials (rates of change in firing rates) that would be predicted under the neural process theory depicted in figure 2 (Friston et al., 2017a). As can be seen, the neuronal populations encoding sadness and panic quickly acquire elevated firing rates, and continue to fire at equivalent rates throughout the rest of the trial (reflecting the belief that both emotion categories are equally probable).

We did note two particular dependencies on other parameters, however. First, the agent’s tendency to “play it safe” and only report the coarse-grained category depended on the aversion to incorrect feedback. As this negative preference was lowered, the agent had stronger and stronger tendencies to guess on each trial, leading to chance levels of accuracy (see figure 8 for an illustration of an example trial). This mimicked what has been observed in childhood during emotion concept learning, where children tend to first use specific emotion terms in a non-specific way (i.e., they are used more or less interchangeably to mean “bad”; (Widen and Russell, 2008)). This also highlights the fact that a person ‘needs to care’ to a sufficient degree about being accurate when conveying their emotions to others. As a further effect, we noted that as the level of emotion concept precision decreased, the magnitude of prior expectations favoring somatic conditions necessary to promote somatization also became lower. Thus, poor emotion concept acquisition also created a greater vulnerability to somatic misattribution.

**Figure 8.**
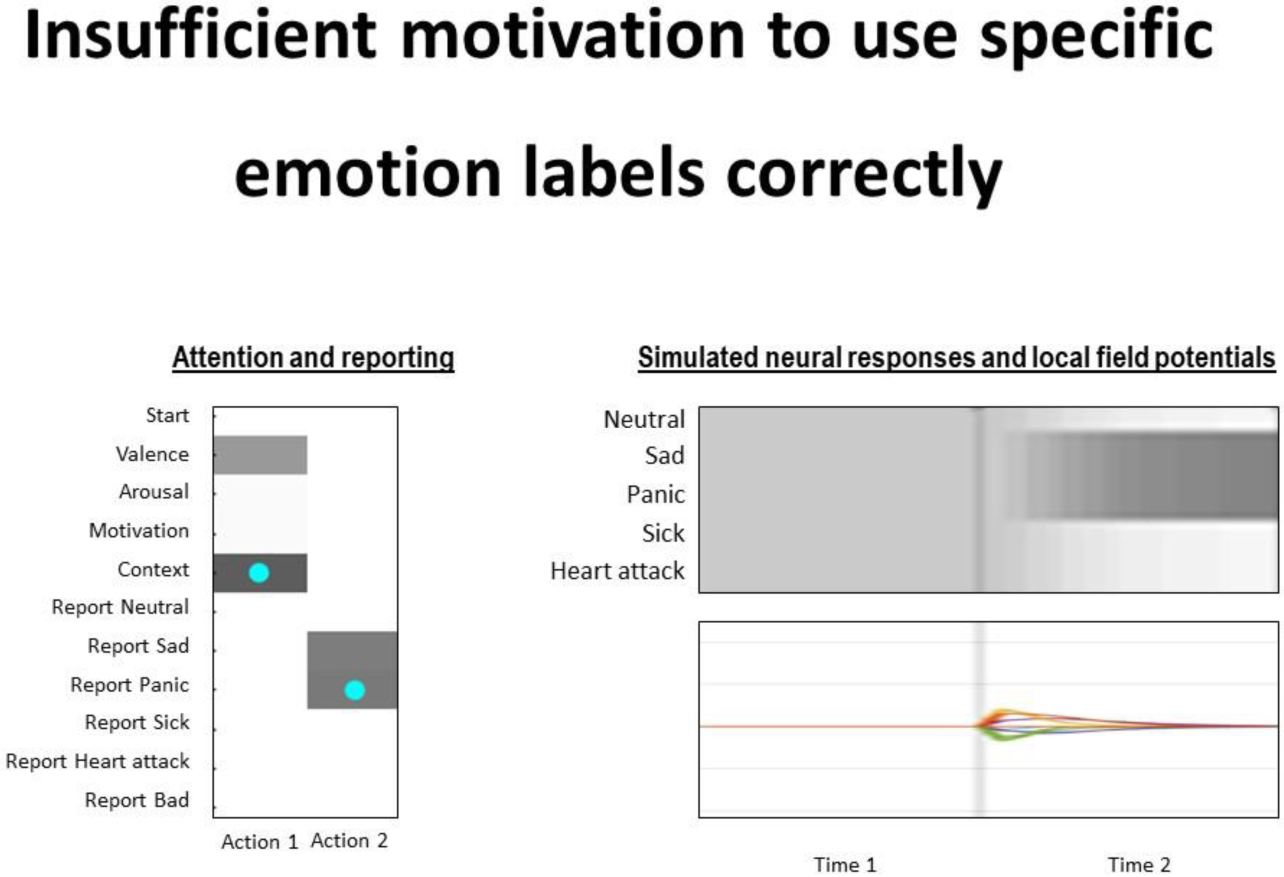
Example trial displaying suboptimal performance as a result of a combination of poor emotion concept acquisition (as in the previous simulation) and an attenuated aversion to reporting incorrectly. As shown on the left, the agent first chose to attend to her beliefs about context (and observed social threat), at which point, despite being equally confident in reporting sad or panic, simply chose to guess one of the two (cyan dots indicate the true action taken; darker colors indicate higher levels of confidence in one action over others). The right panel illustrates the simulated neuronal firing rates (darker = higher firing rate) and local field potentials (rates of change in firing rates) that would be predicted under the neural process theory depicted in figure 2 (Friston et al., 2017a). As can be seen, the neuronal populations encoding sadness and panic quickly acquire elevated firing rates, but neither population outcompetes the other before the agent makes a choice.

### 4.5 Mechanism 4: Biased attention

A fourth mechanism discussed in previous literature pertains to selective attention biases (Lane et al., 2018; Smith et al., 2018a; Smith and Lane, 2016). That is, even if an individual has appropriate prior expectations – and has acquired precise emotion concept knowledge – the maladaptive allocation of selective attention could still hinder evidence accumulation necessary to correctly infer upon one’s own internal states. Here, we simulated two examples of such an attentional bias, by equipping the model’s **E**-matrix (a prior bias over policy selection) with strong prior expectations favoring the selection of some policies over others (see figure 9 for an example trial). We first explored the consequences of equipping the model with the strong habit of focusing on its own bodily state (valence, arousal, and action tendencies), while ignoring external contextual information. Here, we observed that, if the agent possessed these suboptimal attentional or epistemic habits, this promoted extended periods of attentional sampling and self-reported internal states became inconsistent. In other words, the agent could not differentiate between emotional and somatic causes, and therefore simply guessed on each trial between sadness/sickness and panic/heart attack (leading to chance levels of accuracy). This is consistent with previous studies of context effects in emotion recognition, illustrating, for example, that attention to facial information alone is insufficient to infer the emotional states of others, and that available contextual information is necessary to disambiguate between different possible emotions (Aviezer et al., 2008; Barrett et al., 2011). If biased expectations favoring somatic causes were also present under these conditions, we noted that this again promoted somatic misattribution, whereas reductions in emotion concept precision led to greater numbers of coarse-grained emotion reports. Thus, this type of attentional bias could produce confusion/uncertainty about emotional versus somatic causes, but consistent somatic misattribution or coarse-grained reporting still required the further influence of biased prior expectations or poor emotion concept acquisition.

**Figure 9.**
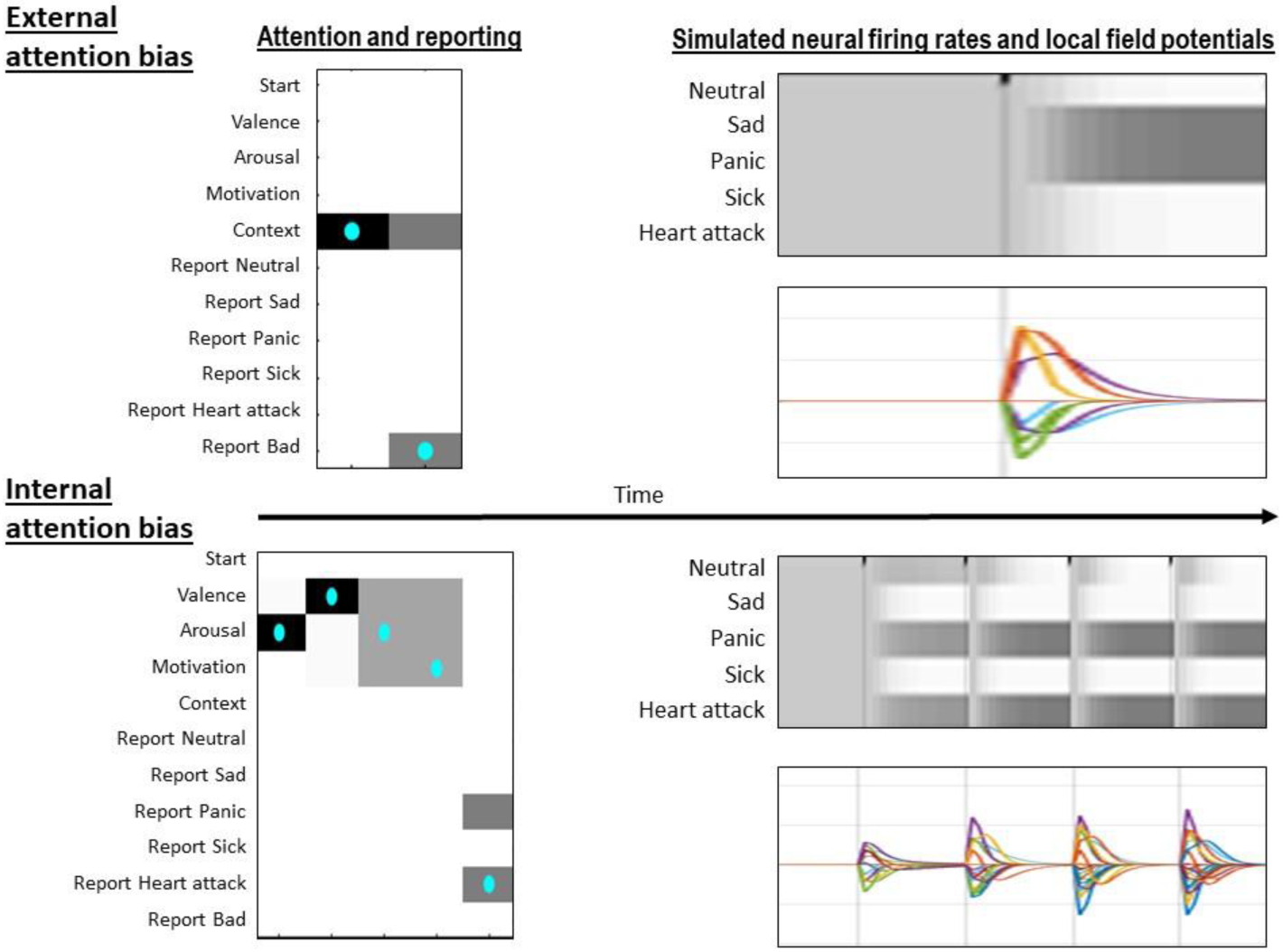
Example trial displaying suboptimal performance as a result of two different types of attentional biases. The top panel illustrates the agent’s behavior in the context of an “external” bias in which she had a strong tendency to focus on the context and ignore bodily sensations. As shown on the top left, this bias only allowed the agent to disambiguate situations that were more or less likely to involve emotions but did not allow her to distinguish finer-grained categories. In this case, as the context involved social threat, she therefore simply reported feeling bad (cyan dots indicate the true action taken; darker colors indicate higher levels of confidence in one action over others). The top right panel illustrates the simulated neuronal firing rates (darker = higher firing rate) and local field potentials (rates of change in firing rates) that would be predicted under the neural process theory depicted in figure 2 (Friston et al., 2017a). As can be seen, the neural populations encoding sadness and panic quickly acquire elevated firing rates, but neither population outcompetes the other before the agent makes a choice. The bottom panel illustrates the agent’s behavior in the context of an “internal” bias in which she had a strong tendency to focus on bodily sensations and ignore context. This resulted in a greater number of attentional shifts (i.e., slower reaction time) before responding, and precluded the ability to distinguish between emotional and non-emotional bodily states – on this trial, leading her to guess heart attack after observing high arousal, negative valence, and avoidance motivation. The associated simulated neuronal firing rates and local field potentials reflect an initial decrease in confidence in the states of sadness and sickness, and then subsequent decreases in confidence in a neutral state and simultaneous (equivalent) increases in confidence in both panic and heart attack over time.

We subsequently simulated an external attentional bias where the agent ignored its affective bodily responses and only attended to contextual information. In this case, on emotion trials it reported that it felt “bad” in 100% of cases. Thus, an external bias could also promote behavior consistent with coarse-grained, “level 2” emotional awareness. This external bias bears some conceptual similarity to constructs currently measured via self-report scales of low emotional awareness (e.g., the externally oriented thinking tendency measured by the Toronto alexithymia scale [TAS-20]; (Bagby et al., 1994b)).

### 4.6 Mechanism 5: Beliefs favoring high emotional volatility

A fifth mechanism that could produce reductions in emotional awareness corresponds to the belief that emotional states are highly unstable. Of course, an individual’s affective responses could in fact be highly volatile, or these beliefs could be exaggerated; borderline personality disorder, for example, is characterized by highly volatile emotions, and has also been associated with lower levels of emotional awareness (Levine et al., 1997). To illustrate the effects of this mechanism, we applied the same softmax manipulation described above (precision = 0.01) to the model’s **B**-matrices that specified beliefs about the probability that each emotional state would transition to a different internal state across the trial, but kept the actual affective response features stable across the trial. This caused the agent to repeatedly attend to the same information over and over again (to repeatedly check whether the state had changed; similar to the simulation shown in figure 7) and then simply report that she felt bad (85-100% of trials) or occasionally report a somatic misattribution (i.e., believing that she may have started out in an emotional state but that she was now in a dangerous somatic state; e.g., “This may have started out as a panic attack, but now I’m definitely having a heart attack.”).

### 4.7 Summary of first-level simulation results

In summary, these simulations highlight a range of interacting mechanisms that can each produce (either in isolation or combination) the somatic misattribution and low emotion granularity phenomena characteristic of the lower levels of trait emotional awareness. Each mechanism or combination produced different profiles, involving either somatic misattribution, coarse-grained emotion reports, or the inconsistent use of specific emotion terms. Each was also associated with different amounts of time (i.e. number of attentional shifts) before the agent chose to make its report, which could be understood as different reaction times in a behavioral task. This suggests that – for an individual to display higher levels of emotional awareness, in which specific emotion concepts can be adaptively and reliably used to understand their own affective responses – none of these mechanisms can interfere with emotion-related belief updating. *That is, affective response generation mechanisms must be functioning adaptively, a person must expect that emotions are likely to occur, they need to have acquired precise emotion concept knowledge, they must possess attentional habits that incorporate both bodily and contextual factors in the inference process, and they need to believe that emotional states are sufficiently stable to infer them by accumulating evidence over time*.

### 4.8 Mechanism 6: Access to domain-general cognition

While we allowed for first-level reporting behavior in the simulations described above, this was primarily to illustrate the effect of confidence and biases on the granularity of emotion reports. As noted above, in the three-process model (Smith et al., 2018a), and other neuro-cognitive models of conscious awareness more generally (e.g., global neuronal workspace models; (Dehaene, 2014)), internally representing information is not sufficient for reportable awareness (e.g., studies have shown that sensory stimuli – including those associated with emotion concepts – can produce both perceptual and conceptual/semantic priming effects on behavior in the absence of self-reported experience/recognition of those stimuli, and top-down attention can also have influences outside of awareness; (Dehaene et al., 2014, 2006; Panksepp et al., 2017; Schnuerch et al., 2016; Smith, 2019; Smith and Lane, 2016; Zemack-Rugar et al., 2007)). These models suggest that, in addition, reportable awareness also requires that the perceptual/conceptual contents of a given neural representation (i.e., the electrochemically encoded messages that it passes to other neuronal populations) are selectively “gated” or “broadcast” to higher-order cognitive systems that have limited capacity (i.e., these higher-order cognitive systems are suggested to correspond to distributed, large-scale neural networks spanning medial and lateral frontoparietal association cortices). Thus, in addition to inferring one’s own most probable internal state as we simulated above, a model of emotional awareness should also include this type of gating mechanism, and a failure of emotional state information to be made accessible to higher-order cognition could represent an additional factor promoting low emotional awareness.

To simulate this type of mechanism, we enabled the second hierarchical level of our model associated with emotion-focused working memory. We then again used a softmax function (inverse temperature parameter = 0.01) to manipulate the precision of the mapping from working memory contents to lower-level internal state representations – as, in active inference, this would determine the degree to which emotion concepts are reliably broadcast into working memory. In other words, assigning higher or lower levels of precision to a given representation (i.e., which could be done in a context-specific or goal-specific manner) determines whether or not that representation has a significant influence in updating the contents of domain-general cognition. When manipulating the precision of the mapping between particular emotion concepts and working memory in our model, we were able to confirm this expected effect. That is, when the precision of this mapping was lowered, emotion concepts did not gain appropriate access to and update working memory contents, leading to chance accuracy on the working memory task.

Figure 10 illustrates an exemplar case, where the precision of the mapping from sadness and panic into working memory was reduced. This mimics an individual who, while having a good conceptual understanding of such emotions, and perhaps reacting in a manner consistent with such emotions, may ignore the possibility that they are feeling emotions and focus on other types of information. For example, someone who believes that focusing on certain emotions is a sign of weakness or otherwise considers this type of information of low value. In the example trial shown, the posterior probability distribution representing working memory contents remains highly imprecise, while the lower-level emotional state representations remain precise (i.e., a potential example of unconsciously represented emotion). The agent also was incapable of reflecting on her emotions and correctly determining whether her emotion at the third time point (neutral) was similar to her earlier emotions. Across 60 trials (10 trials for each of the six internal state combinations that could be held in working memory), we confirmed that low levels of precision for lower-level emotions rendered the agent capable of accessing and holding other somatic internal states in working memory – and performing the comparison task correctly with respect to other somatic states (i.e., the state corresponding to sickness and panic attack) – but failed to reliably access or use working memory contents that included the concepts of sadness and/or panic. In short, reportable awareness of single emotions or emotion combinations (characteristic of “level 4” emotional awareness) were compromised, while other internal state representations remained accessible.

**Figure 10.**
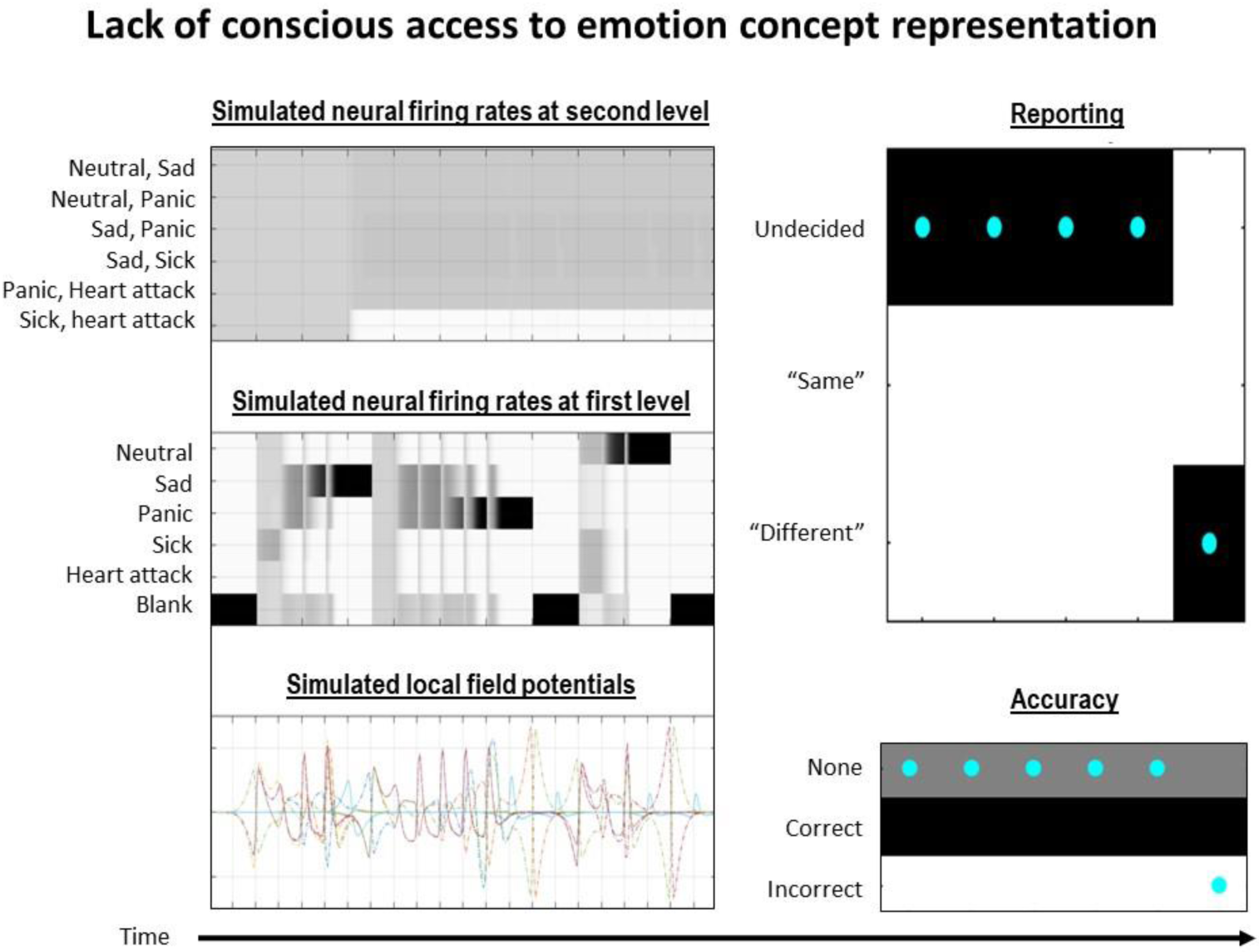
Displays suboptimal inference of a combination of two emotions in the higher-level model as a result of reduced precision in the mapping between represented emotional states at the first level and working memory contents at the second level – leading to a failure to “broadcast” represented emotion concepts into domain-general cognition. Shown are the simulated neuronal firing rates and local field potentials generated by both the first and second levels of the model. As can be seen, at the first level the first emotion represented was sadness and the second was panic. The third represented emotion was neutral. In contrast, it can be seen that the higher-level is not updated to include information about sadness and panic (i.e., it only knows that the state isn’t sickness and heart attack), preventing reportable emotional awareness and successful task performance (right).

### 4.9 Mechanism 7: Working memory content stability

The final mechanism we considered corresponds to previous empirical observations that higher emotional awareness is associated with greater emotion-focused working memory capacity (R Smith et al., 2017b). As shown in previous simulation work (Parr and Friston, 2017b), once information has been gated into working memory, the ability to maintain and manipulate working memory contents over a delay period then depends on the rate with which that information decays – which in turn depends on the stability of the contents of working memory, here operationalized as the precision of the model’s **B**-matrix for the hidden state factor corresponding to working memory contents. In other words, working memory capacity will be reduced (i.e., information will decay more quickly) if past and future states are assumed to be less predictable from present states. Thus, the greater the probability that working memory contents will defuse away from their current state, the more difficult it will be to maintain and reflect on multiple emotions. This is a necessary condition for the highest levels of emotional awareness (i.e., in which an individual can contemplate multiple emotions at once, including both their own emotions and someone else’s).

To simulate differences in emotion-focused working memory capacity, we therefore manipulated the precision of the model’s transition beliefs about working memory contents using the same manipulation as in previous simulations applied to the relevant **B**-matrix (precision = 0.5). Figure 11 illustrates the results of an exemplar trial. As can be seen, the posterior probability distribution over hidden states of the second level disperses over time until it becomes highly imprecise, corresponding to an inability to maintain precise contents. This was confirmed in 100 repeated simulations

**Figure 11.**
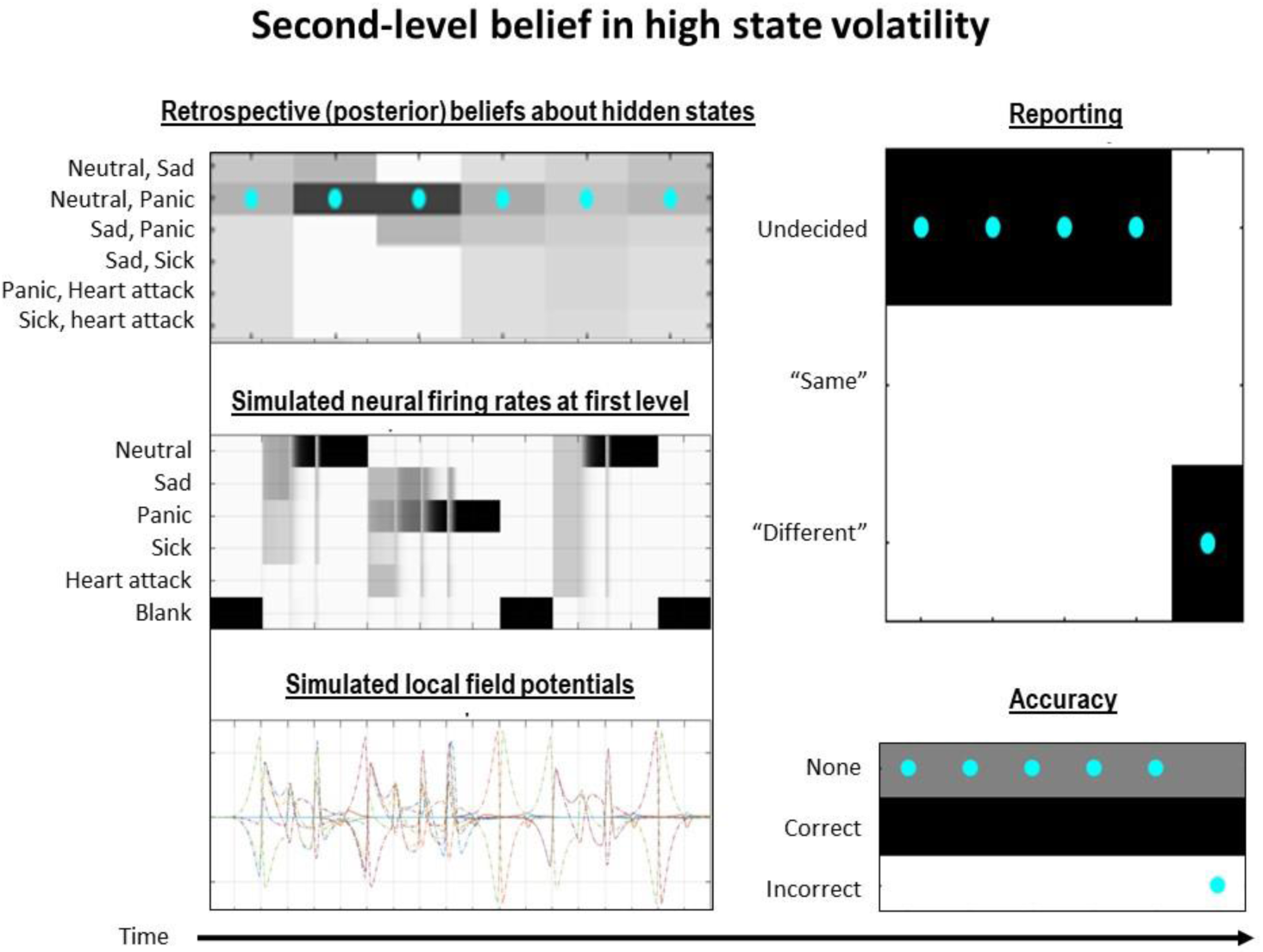
Displays an inability of the higher-level model to hold lower-level information in working memory as a result of reduced transition precision, such that the mapping between the contents of working memory at one point in time does not precisely predict the contents of working memory at past and future times. Shown on the top left are the agent’s posterior (retrospective) beliefs regarding higher-level states across the trial. As can be seen, while the contents associated with the true state (cyan dots) are precise at the second and third time point, there is subsequent decay such that, by the end of the trial, beliefs over states become highly imprecise – leading to poor task performance. Also shown are the simulated first-level firing rates, and the local field potentials generated by both the first and second levels of the model. As discussed in the text, low transition precision could reflect a stable trait difference, but it could also reflect temporary reductions in internal estimates of the reliability of long timescale regularities during stressful or otherwise high arousal situations.

In individuals with low emotional awareness, reductions in high-level transition precision could plausibly operate in both a state and trait manner. For example, emotional awareness could be constrained in a trait manner, as there appear to be stable individual differences in working memory capacity generally – and in emotion-focused working memory specifically (Melby-Lervåg and Hulme, 2013; R Smith et al., 2017b). In contrast, it is also plausible that estimates of transition precision over longer timescales – associated with working memory – could be modulated in a context-dependent manner. For example, it may be that under threatening or otherwise stressful conditions, information about longer timescales is implicitly estimated as less informative/precise, as such situations may require immediate reflexive action (e.g., evolutionarily, extended deliberation in such cases may have led to death). This would correspond to an attenuation of transition precision, which could temporarily reduce working memory capacity as simulated here. This corresponds to a large body of previous work that has demonstrated reductions in reflective capacity – and reductions in prefrontal neural firing rates that would be predicted by our model – under conditions of high stress and autonomic arousal (Arnsten, 2015; Teigen, 1994). It is also consistent with previous empirical results indicating that emotional awareness levels can fluctuate in a state-dependent manner (Versluis et al., 2018).

## 5. Clinical and Research Implications

In this paper we have used simulations – based on a computational implementation of the three-process model (Smith et al., 2018a) – to demonstrate quantitatively distinct mechanisms that could contribute to individual differences in emotional awareness. These simulations highlight how a single, clinically relevant psychological construct, measured via self-report behavior, need not have a 1-to-1 correspondence to a single aberrant process. Instead, at the level of information processing simulated here, multiple processes could contribute, either in isolation or in combination, to produce the same measured phenotype. This could be important, as low emotional awareness (or the related construct of alexithymia) is a notoriously difficult condition to treat (Ogrodniczuk et al., 2011). One way our model might help is by providing a framework that allows clinicians to systematically consider each of the seven mechanisms illustrated above as possible targets for therapeutic intervention. Further, as reviewed in the introduction, reduced emotional awareness manifests in a large number of conditions (e.g., somatic symptom disorders, depression, substance dependence, autistic spectrum disorders, schizophrenia, PTSD, eating disorders). However, the basis for reduced emotional awareness in each of these clinical conditions has not been systematically studied. Our model could also potentially facilitate future work in this area, by providing the basis for testable hypotheses (e.g., perhaps different underlying mechanisms are at work in different disorders).

For some examples of ways in which mechanistic knowledge could guide the development and selection of individualized treatment, see table 1. In this table, we list each of the seven mechanisms introduced above and highlight potential ways in which they might relate to assessment and treatment. Established treatment modalities that address the mechanisms in question are listed, although it is acknowledged that each modality listed is actually more complex and broader in scope than the mechanisms listed. The table also implicitly highlights potential gaps where new assessment tools and interventions could be useful. For example, if the primary contributing mechanism involved maladaptive affective response generation, effective therapeutic interventions might attempt to modify the way individuals evaluate the situations they perceive and represent, as in current cognitive therapies (Barlow et al., 2016), such that more adaptive automatic responses are generated. If, in contrast, the major contributing mechanism involves maladaptively strong prior expectations for dangerous somatic conditions, the relevant intervention point would likely involve addressing and correcting such expectations. For example, if a therapist facilitates the repeated experience and recognition of emotions during (and outside of) the therapeutic setting, this could plausibly increase a client’s prior expectations for emotions – increasing the chances that future affective responses will be interpreted as emotions instead of as somatic conditions, as in emotion-focused therapy (Greenberg, 2010). Given our observation that, in our particular implementation, a 7-fold greater expectation for a somatic interpretation was associated with reduced emotional awareness, a clinical hypothesis to be tested is that reducing but not eliminating this bias, say reducing it by half, could have significant clinical benefits.

**Table 1.** Assessments, measures, and interventions potentially relevant to each of the computational mechanisms discussed.

| Computational mechanism | Clinical Assessment | Relevant measures | Relevant interventions |
| --- | --- | --- | --- |
| Maladaptive affective response generation | 1. Evaluate for flat or exaggerated affect and situational appraisal/interpretation biases | 1. Peripheral physiological measures<br>2. Valence/arousal rating measures<br>3. Appraisal bias measures<br>(Lang et al., 2008; Scherer, 2009; Scherer and Brosch, 2009; Siemer et al., 2007; Smith et al., 2019b) | 1. Beta blocker to attenuate high arousal states<br>2. Cognitive therapeutic interventions involving identifying automatic situational interpretations and alternative interpretations<br>3. Exposure therapy<br>4. Acceptance and commitment therapy<br>5. Emotion Regulation Therapy<br>(Barlow et al., 2016; Foa et al., 2007; Hayes and Smith, 2005; Mennin and Fresco, 2014) |
| Somatically biased prior expectations | 1. Evaluate for beliefs about the likelihood of somatic threats and past experience of trauma and illness | 1. Anxiety sensitivity index<br>(Mueller and Alpers, 2006) | 1. Psychoeducation about bodily expression of emotions and the benign nature of most somatic sensations<br>2. Focusing – deriving emotional meaning from bodily sensations in context<br>(Gendlin, 1982; Lumley et al., 2017) |
| Poor emotion concept acquisition | 1. Evaluate a client's self-reported beliefs about the causes of specific emotions, and the typical thoughts, bodily feelings, and behaviors associated with those emotions | 1. TAS-20: difficulty describing and identifying feelings subscales (Factors 1 and 2)<br>2. Emotion understanding subscales within trait and ability emotional intelligence tests<br>3. Emotion granularity measures<br>(Kashdan et al., 2015; Lane et al., 1990; Mayer et al., 2003; Parker et al., 2003; Petrides et al., 2016) | 1. Psychoeducation—teaching emotion concepts<br>2. Practice identifying one's own emotions, their causes, and the bodily sensations and behaviors that typically follow<br>3. Emotion-focused therapy<br>(Greenberg, 2010; Hayes and Smith, 2005) |
| Biased attention | 1. Evaluate for tendencies to primarily focus externally or primarily focus only on bodily sensations | 1. TAS-20: externally oriented thinking subscale (Factor 3)<br>(Parker et al., 2003) | 1. Attention bias modification<br>2. Mindfulness/meditation training (Mindfulness-Based Stress Reduction)<br>(Segal et al., 2004) |
| High emotional volatility | 1. Evaluate for mood stability | 1. Personality measures of neuroticism<br>(Costa and McCrae, 1992) | 1. Training adaptive emotion regulation habits (e.g., reappraisal, acceptance, etc.)<br>2. Dialectical Behavior Therapy<br>3. Emotion Regulation Therapy<br>(Mennin and Fresco, 2014; Swales, Heidi L. Heard, J. Mark G., 2000) |
| Conscious inaccessibility | 1. Evaluate attitudes about emotions (e.g., do they provide useful information?; Are they a sign of weakness?) | 1. Measures of cognitive suppression tendencies in emotion regulation | 1. Psychoeducation with regard to the value of emotions<br>2. Correcting emotion avoidance tendencies |
|  |  | (Gross, 1998a; Gross and Levenson, 1997; Spaapen et al., 2014) | 3. Psychodynamic psychotherapy (analysis of defense)<br>(Barlow et al., 2016) (Brenner, 1973) |
| Reduced reflective capacity | 1. Evaluate whether problematic behaviors in a client's life tend to occur in high arousal situations<br>2. Evaluate whether problems pertain to impulsivity or insufficient reflection<br>3. Evaluate for general cognitive ability | 1. Measures of future-oriented thinking<br>2. Measures of emotion-focused working memory capacity<br>(R Smith et al., 2017b; M. Toplak et al., 2014; M. E. Toplak et al., 2014) | 1. Training adaptive emotion regulation habits<br>2. Interventions designed to counter impulsive behavior<br>3. Mentalization-based therapy<br>(Fonagy and Luyten, 2009) |

Next consider an individual whose primary contributing mechanism involves poor emotion concept acquisition. In this case, the most sensible intervention would plausibly involve psychoeducation (Burum and Goldfried, 2007; Lumley et al., 2017). That is, an individual would need to be given the opportunity to gain greater conceptual understanding of the content of the different emotion categories employed within their particular culture. In contrast, if an individual’s primary issue involved biased attention, cognitive-behavioral interventions – in which an individual explicitly practices and keeps records of the thoughts, feelings, and action tendencies they experience in particular situations – would likely be relevant to countering such maladaptive attentional habits, as would more recent mindfulness-based approaches (Barlow et al., 2016; Hayes and Smith, 2005; van der Velden et al., 2015). Of course, many individuals may have multiple contributing mechanisms in play, and these mechanisms can interact in significant ways. For example, if an individual has high prior expectations for somatic conditions, our simulations suggest that they will be more likely to selectively attend to information that would confirm these expectations if they were correct, and then jump to a conclusion before attending to other informative signals. The therapeutic task in such cases would clearly be more complex, corresponding to the added difficulty of identifying and targeting multiple simultaneous and mutually reinforcing mechanisms.

Moving on to consider higher-level cognitive factors, if an individual tends to ignore or avoid reflecting on their own emotional state – here operationalized as failing to gate emotion concept representations into working memory – one major therapeutic task would likely involve identifying the underlying driver of this tendency. For example, if an individual’s past experience has not afforded or motivated considerable attention to emotions, this could lead simultaneously to poor emotion concept acquisition and to a higher-level estimate that emotion concept representations convey unreliable information to higher processing levels (for simulation work illustrating the role of biased attention in preventing emotion concept learning, see (Smith et al., 2019d)). A somewhat (but not completely) overlapping possibility, might be that an individual has developed the habit of ignoring thoughts about emotions for value-based reasons, such as the belief that information about emotions is not useful in decision-making or that reflecting on emotions entails a type of weakness or undesired vulnerability (e.g., as in cases of gender-and culture-based socialization; (Chaplin et al., 2005)). Optimal therapeutic interventions could then presumably target such beliefs through psychoeducation regarding the value/usefulness of emotions. Yet another focus could be sources of insecurity or fear of vulnerability that are driving cognitive strategies aimed at emotion avoidance, or the related fear that thinking/talking about previous emotionally troubling experiences could generate more intense discomfort (e.g., as in individuals with PTSD (Foa et al., 2007)).

The final mechanism we have considered involves reduced higher-level transition precision, which would promote reduced working memory capacity. It is currently unclear whether this type of trait difference in transition precision is malleable, and previous attempts to improve working memory capacity have met with limited success (Melby-Lervåg and Hulme, 2013). One interesting possibility that should be investigated in future research is whether successful interventions could be designed that would allow individuals to learn that long timescale regularities are more reliable. For example, individuals who grow up in stressful and unpredictable environments appear to learn that distant future states/outcomes are unpredictable, leading decision-making to focus on achieving proximal versus distal goals (e.g., steeper delay discounting, greater risk-taking, etc.; (Kavanagh and Kahl, 2018)). If such individuals could learn that distant future states are more predictable in their current adult environment, this could potentially promote greater reflective tendencies (i.e., it would be more internally rational to “bet on” predictions about the distant future when making decisions). However, there is insufficient evidence at present to assess the plausibility of this possibility.

On the other hand, state differences in the transition precision of working memory contents may be more therapeutically addressable. Specifically, consider cases where recurring context-dependent influences, such as high levels of stress and autonomic arousal, lead to repeated situations in which a person’s reflective capacity is reduced (e.g., leading to impulsive and suboptimal decision-making). For example, if this occurs as a result of chronic stress, then this could be addressed by practicing emotion regulation strategies (e.g., as in dialectical behavior therapy or emotion regulation therapy; (Mennin and Fresco, 2014; Swales, Heidi L. Heard, J. Mark G., 2000)) or finding ways of preventing/improving recurring stressful or problematic situations. In some cases, exposure therapies could also be beneficial if they help a person learn that they can handle being in a fear-provoking situation; it could also potentially lead to reductions in this type of situation-dependent arousal and therefore improve reflective capacity in such contexts. One would expect that if an individual is better able to understand and reflect on how they’re feeling before acting in such situations that their decision-making would become more adaptive.

The above considerations, while speculative, illustrate the way that an active inference formulation of emotion-related processes – even the simple toy model presented here – may be able to further clinical thinking. In addition, it could also guide complementary empirical research. As we have shown, under the neural process theory associated with active inference (Friston et al., 2017a), many of these mechanisms are predicted to produce different patterns of neural firing rates and local field potentials that could be tested in neuroimaging paradigms. Some mechanisms should also be associated with faster responding times than others, depending on whether they promote overconfidence and “jumping to conclusions” or instead promote continued information gathering behavior as a result of low confidence. In principle, this suggests that different reaction times when people are asked to report their emotions could provide evidence for the operation of one mechanism versus another. Given an appropriate emotion reporting task, neuroimaging studies could also test for the predicted patterns of neural responses in specific brain regions (e.g., activation of default mode network regions associated with emotion conceptualization processes, or executive control network regions associated with higher-level working memory processes; (Barrett and Satpute, 2013; Binder et al., 2009; Rottschy et al., 2012; Seeley et al., 2007)).

It is important to stress the simplified nature of the model we have presented. For example, while we were able to simulate the ambiguous, probabilistic mapping between emotion concepts and lower-level perceptual experiences such as the valence, arousal, and action tendencies felt in one’s body during affective responses (i.e., consistent with constructivist accounts of emotion conceptualization; (Barrett, 2017)), these were modeled purely as binary variables, whereas in reality they involve multiple levels and dimensions (Colibazzi et al., 2010; Posner et al., 2005). This suggests that emotion conceptualization likely draws on a much richer generative model, with more complex mappings to lower-level representations. Furthermore, most people also have many more emotion concepts and somatic concepts than we considered. That said, the general mechanisms we have simulated – with regard to the precision of concept-percept mappings, attentional biases, prior expectations, and beliefs about the predictability of future states – would still be expected to hold in a more complete generative model. Self-directed emotion recognition tasks that would allow behavior to be fit to our model will need to be designed to confirm this possibility.

There are some additional future directions that could be considered in relation to our model. First, although we have illustrated how neurocomputational processes can afford simple cognitive or behavioral uses of emotional state inference, the adaptive use of emotional awareness in guiding more complex behavior was not modeled. This represents an important direction for future work. As one example, an important ability that is plausibly facilitated by emotional awareness is the broader construct of emotion regulation (ER; (Aldao, 2013; Morrish et al., 2019)) – which includes many subtypes, such as taking action to modify emotion-provoking situations, engaging in cognitive strategies to adjust one’s interpretation of those situations, and suppressing maladaptive automatic action tendencies (Gross, 1998b). Many of these strategies can be understood in terms of interactions between conscious access processes and affective response generation processes within the TPM (e.g., manipulating situational interpretations in working memory, so as to alter subsequent affective response generation). Understanding how one is feeling (i.e., emotional state representation) plausibly represents an important piece of information that could adaptively guide the use of each of these strategies (e.g., in guiding attention to the cognitive or situational sources of that emotion, informing predictions about its time course given different actions, etc.); extending our model to include a policy space of voluntary emotion regulation strategies could therefore provide additional mechanistic insights of potential clinical relevance (Aldao et al., 2010). There is also evidence that emotional awareness can facilitate a kind of “automatic” emotion regulation, in which simply identifying how one is feeling can lead to reduced emotional arousal (Kircanski et al., 2012). This opens up an additional possibility for future work to model the way that top-down influences of emotional state inference can directly modulate lower-level visceral policy selection. Indeed, one could argue that emotion regulation implies the existence of higher-level egocentric representations that entail a minimal selfhood. These would be necessary to issue descending predictions of precision that constitute internal policies (i.e., mental actions) that – on an active inference reading of emotion processing (Badcock et al., 2017; Fotopoulou and Tsakiris, 2017; Seth and Friston, 2016) – might constitute emotion regulation.

Yet another opportunity to extend our model in future work would be to incorporate the influence of other higher-level cognitive biases (e.g., expectations of low reward, negativity biases, etc.) on emotion inference and somatovisceral policy selection, such as those often included in cognitive and computational models of depression and that have been linked to particular neural systems (e.g., dopaminergic and serotonergic dysfunction in depression; (Adams et al., 2015; Chekroud, 2015; R Smith et al., 2017a)). While we did model a form of top-down attentional bias, our model did not include the types of depressive schema-based interpretation biases that are also of significant clinical importance – and that could also interact with emotional awareness in interesting ways. For example, negativity biases could in part be associated with (and modeled as) precise prior expectations to experience negatively valenced states, which could in turn bias emotional state inference (and emotion concept learning during development; see (Smith et al., 2019d)) away from recognition of neutral or positive emotions.

Finally, future modeling work should also incorporate the role of bidirectional interactions with the environment as they pertain to the generation and internal representation of emotions. In active inference models, this can be characterized by separately simulating the generative process (the actual environmental dynamics giving rise to an agent’s sensations) and the generative model (the agent’s beliefs about the causes of those sensations). In the present context, for example, this could afford modeling the types of social interactions with other agents in which more or less adaptive affective responses could be generated by different inferences about others’ emotional states, and in which iterative interactions with other agents could be guided more or less adaptively by different inferences about one’s own emotional states. Such an approach could fruitfully build on recent work using active inference to model social and environmental niche construction (Bruineberg et al., 2018; Constant et al., 2018).

## 6. Conclusion

To conclude, we have used the example of trait emotional awareness to illustrate the way that clinically relevant individual differences can be produced by a range of underlying neuro-computational mechanisms (for related work on computational mechanisms of psychotherapy, see (Moutoussis et al., 2017; Smith et al., 2019c)). Our simulations suggest that many underlying processes can combine in different ways to produce the same observable clinical phenomenon. Formally, this speaks to a degenerate (many-to-one) mapping between pathophysiology and psychopathology. In principle, this should also apply to other clinical phenomena that are measured via self-reported experience. This highlights the need to identify the underlying processes at work in any given individual and to design/implement interventions targeting those particular processes in those individuals. Our hope is that this model can inspire neuroimaging and behavioral paradigms that, in conjunction with this type of model, could help in identifying these mechanisms and eventually inform treatment selection.

## Software note

Although the generative model – specified by the various matrices described in this paper – changes from application to application, the belief updates are generic and can be implemented using standard routines (here **spm_MDP_VB_X.m**). These routines are available as Matlab code in the SPM academic software: http://www.fil.ion.ucl.ac.uk/spm/. The simulations in this paper can be reproduced (and customised) via running the Matlab code included here is supplementary material (**MDP_EA_final.m**).

## Footnotes

1 The term “conscious access” is taken from global neuronal workspace (GNW) models of conscious awareness (Dehaene et al., 2014), which are based on a body of empirical work examining neural activity underlying the reportable experience of sensory stimuli generally (for a review, see (Dehaene, 2014)). According to these models, reportable conscious experience requires that perceptual information is made widely accessible (through a type of selective “broadcasting” process) to the large-scale frontal-parietal networks that underlie domain-general cognition. While necessary for *reportable* awareness, it remains an open debate whether this type of “access” is necessary for phenomenal experience (“what it is like”) itself. Some authors argue that it is necessary (e.g., see (Dehaene, 2014; Smith, 2017, 2016)), whereas others instead argue that an individual could subjectively experience a stimulus while concurrently self-reporting no awareness of having seen (heard, felt, etc.) that stimulus (e.g., see (Block, 2005)). The TPM, as described here, does not commit to a position in this debate. As used here, “conscious access” refers only to those processes necessary for self-reportable awareness of one’s emotions and the use of that information in guiding goal-directed decision-making.

## Notes

#### Summary of Updates

Additional figure added (figure 2). Added paragraphs to the discussion on emotion regulation.

## References

Adams, R.A., Huys, Q.J.M., Roiser, J.P., 2015. Computational Psychiatry: towards a mathematically informed understanding of mental illness. J. Neurol. Neurosurg. Psychiatry jnnp-2015–310737. https://doi.org/10.1136/jnnp-2015-310737

Aldao, A., 2013. The Future of Emotion Regulation Research. Perspect. Psychol. Sci. 8, 155–172. https://doi.org/10.1177/1745691612459518

Aldao, A., Nolen-Hoeksema, S., Schweizer, S., 2010. Emotion-regulation strategies across psychopathology: A meta-analytic review. Clin. Psychol. Rev. 30, 217–237. https://doi.org/10.1016/J.CPR.2009.11.004

Allen, M., Friston, K.J., 2018. From cognitivism to autopoiesis: towards a computational framework for the embodied mind. Synthese 195, 2459–2482. https://doi.org/10.1007/s11229-016-1288-5

Allen, M., Levy, A., Parr, T., Friston, K.J., 2019. In the Body’s Eye: The Computational Anatomy of Interoceptive Inference. bioRxiv 603928. https://doi.org/10.1101/603928

Anderson, M., 2014. After Phrenology: Neural Reuse and the Interactive Brain. MIT Press, Cambridge, MA.

Arnsten, A.F.T., 2015. Stress weakens prefrontal networks: molecular insults to higher cognition. Nat. Neurosci. 18, 1376–85. https://doi.org/10.1038/nn.4087

Aviezer, H., Hassin, R., Ryan, J., Grady, C., Susskind, J., Anderson, A., Moscovitch, M., Bentin, S., 2008. Angry, Disgusted, or Afraid? Psychol. Sci. 19, 724–732. https://doi.org/10.1111/j.1467-9280.2008.02148.x

Badcock, P.B., Davey, C.G., Whittle, S., Allen, N.B., Friston, K.J., 2017. The Depressed Brain: An Evolutionary Systems Theory. Trends Cogn. Sci. 21, 182–194. https://doi.org/10.1016/J.TICS.2017.01.005

Bagby, R., Parker, J., Taylor, G., 1994a. The twenty-item Toronto Alexithymia Scale—I. Item selection and cross-validation of the factor structure. J. Psychosom. Res. 38, 23–32.

Bagby, R., Parker, J., Taylor, G., 1994b. The twenty-item Toronto Alexithymia Scale—II. Convergent, discriminant, and concurrent validity. J. Psychosom. Res. 38, 33–40.

Barchard, K., Hakstian, A., 2004. The nature and measurement of emotional intelligence abilities; basic dimensions and their relationships with other cognitive abilities and personality variables. Educ. Psychol. Meas. 64, 437–462.

Barlow, D., Allen, L., Choate, M., 2016. Toward a Unified Treatment for Emotional Disorders - Republished Article. Behav. Ther. 47, 838–853. https://doi.org/10.1016/j.beth.2016.11.005

Barrett, L., 2017. How emotions are made: The secret life of the brain. Houghton Mifflin Harcourt, New York.

Barrett, L., Mesquita, B., Gendron, M., 2011. Context in Emotion Perception. Curr. Dir. Psychol. Sci. 20, 286–290. https://doi.org/10.1177/0963721411422522

Barrett, L., Satpute, A., 2013. Large-scale brain networks in affective and social neuroscience: towards an integrative functional architecture of the brain. Curr. Opin. Neurobiol. 23, 361–72. https://doi.org/10.1016/j.conb.2012.12.012

Barrett, L., Simmons, W., 2015. Interoceptive predictions in the brain. Nat. Rev. Neurosci. 16, 419–29. https://doi.org/10.1038/nrn3950

Baslet, G., Termini, L., Herbener, E., 2009. Deficits in emotional awareness in schizophrenia and their relationship with other measures of functioning. J. Nerv. Ment. Dis. 197, 655.

Berthoz, S., Ouhayoun, B., Parage, N., 2000. Etude preliminaire des niveaux de conscience emotionnelle chez des patients deprimes et des controles. (Preliminary study of the levels of emotional awareness in depressed patients and controls.). Ann. Med. Psychol. 158, 665–672.

Binder, J., Desai, R., Graves, W., Conant, L., 2009. Where Is the Semantic System? A Critical Review and Meta-Analysis of 120 Functional Neuroimaging Studies. Cereb. Cortex 19, 2767–2796. https://doi.org/10.1093/cercor/bhp055

Block, N., 2005. Two neural correlates of consciousness. Trends Cogn. Sci. 9, 46–52. https://doi.org/S1364-6613(04)00318-3 [pii] 10.1016/j.tics.2004.12.006

Bréjard, V., Bonnet, A., Pedinielli, J., 2012. The role of temperament and emotional awareness in risk taking in adolescents. L’Encéphale Rev. Psychiatr. Clin. Biol. Thérapeutique 38, 1–9.

Brenner, C., 1973. An elementary textbook of psychoanalysis. International Universities Press.

Bruineberg, J., Rietveld, E., Parr, T., van Maanen, L., Friston, K.J., 2018. Free-energy minimization in joint agent-environment systems: A niche construction perspective. J. Theor. Biol. 455, 161–178. https://doi.org/10.1016/J.JTBI.2018.07.002

Burum, B.A., Goldfried, M.R., 2007. The Centrality of Emotion to Psychological Change. Clin. Psychol. Sci. Pract. 14, 407–413. https://doi.org/10.1111/j.1468-2850.2007.00100.x

Bydlowski, S., Corcos, M., Jeammet, P., Paterniti, S., Berthoz, S., Laurier, C., Chambry, J., Consoli, S., 2005. Emotion-processing deficits in eating disorders. Int. J. Eat. Disord. 37, 321–329.

Chaplin, T., Cole, P., Zahn-Waxler, C., 2005. Parental Socialization of Emotion Expression: Gender Differences and Relations to Child Adjustment. Emotion 5, 80–88. https://doi.org/10.1037/1528-3542.5.1.80

Chekroud, A.M., 2015. Unifying treatments for depression: an application of the Free Energy Principle. Front. Psychol. 6, 153. https://doi.org/10.3389/fpsyg.2015.00153

Ciarrochi, J., Caputi, P., Mayer, J., 2003. The distinctiveness and utility of a measure of trait emotional awareness. Pers. Individ. Dif. 34, 1477–1490.

Clark, J., Watson, S., Friston, K., 2018. What is mood? A computational perspective. Psychol. Med. 1–8. https://doi.org/10.1017/S0033291718000430

Colibazzi, T., Posner, J., Wang, Z., Gorman, D., Gerber, A., Yu, S., Zhu, H., Kangarlu, A., Duan, Y., Russell, J.A., Peterson, B.S., 2010. Neural systems subserving valence and arousal during the experience of induced emotions. Emotion 10, 377–389. https://doi.org/2010-09991-007 [pii] 10.1037/a0018484

Conant, C., Ashbey, W., 1970. Every good regulator of a system must be a model of that system. Int. J. Syst. Sci. 1, 89–97. https://doi.org/10.1080/00207727008920220

Consoli, S., Lemogne, C., Roch, B., Laurent, S., Plouin, P., Lane, R., 2010. Differences in emotion processing in patients with essential and secondary hypertension. Am. J. Hypertens. 23, 515–521.

Constant, A., Ramstead, M.J.D., Veissière, S.P.L., Campbell, J.O., Friston, K.J., 2018. A variational approach to niche construction. J. R. Soc. Interface 15, 20170685. https://doi.org/10.1098/rsif.2017.0685

Costa, P., McCrae, R., 1992. Revised NEO Personality Inventory (NEOPI-R) and NEO Five-Factor Inventory (NEO-FFI) professional manual. Psychological Assessment Resources, Inc., Odessa, FL.

de Berker, A., Rutledge, R., Mathys, C., Marshall, L., Cross, G., Dolan, R., Bestmann, S., 2016. Computations of uncertainty mediate acute stress responses in humans. Nat. Commun. 7, 10996. https://doi.org/10.1038/ncomms10996

Dehaene, S., 2014. Consciousness and the Brain. Viking Press.

Dehaene, S., Changeux, J., Naccache, L., Sackur, J., Sergent, C., 2006. Conscious, preconscious, and subliminal processing: a testable taxonomy. Trends Cogn. Sci. 10, 204–11. https://doi.org/10.1016/j.tics.2006.03.007

Dehaene, S., Charles, L., King, J.-R., Marti, S., 2014. Toward a computational theory of conscious processing. Curr. Opin. Neurobiol. 25, 76–84. https://doi.org/10.1016/j.conb.2013.12.005

Donges, U., Kersting, A., Dannlowski, U., Lalee-Mentzel, J., Arolt, V., Suslow, T., 2005. Reduced awareness of others’ emotions in unipolar depressed patients. J. Nerv. Ment. Dis. 193, 331–337.

Foa, E., Hembree, E., Rothbaum, B., 2007. Prolonged Exposure Therapy for PTSD: Emotional Processing of Traumatic Experiences Therapist Guide. Oxford University Press.

Fonagy, P., Luyten, P., 2009. A developmental, mentalization-based approach to the understanding and treatment of borderline personality disorder. Dev. Psychopathol. 21, 1355–81. https://doi.org/10.1017/S0954579409990198

Fotopoulou, A., Tsakiris, M., 2017. Mentalizing homeostasis: the social origins of interoceptive inference – replies to Commentaries. Neuropsychoanalysis 19, 71–76. https://doi.org/10.1080/15294145.2017.1307667

Frewen, P., Lane, R., Neufeld, R., Densmore, M., Stevens, T., Lanius, R., 2008. Neural correlates of levels of emotional awareness during trauma script-imagery in posttraumatic stress disorder. Psychosom. Med. 70, 27–31.

Friston, K., FitzGerald, T., Rigoli, F., Schwartenbeck, P., O Doherty, J., Pezzulo, G., 2016. Active inference and learning. Neurosci. Biobehav. Rev. 68, 862–879. https://doi.org/10.1016/j.neubiorev.2016.06.022

Friston, K., FitzGerald, T., Rigoli, F., Schwartenbeck, P., Pezzulo, G., 2017a. Active Inference: A Process Theory. Neural Comput. 29, 1–49. https://doi.org/10.1162/NECO_a_00912

Friston, K., Parr, T., de Vries, B., 2017b. The graphical brain: Belief propagation and active inference. Netw. Neurosci. 1, 381–414. https://doi.org/10.1162/NETN_a_00018

Friston, K., Rosch, R., Parr, T., Price, C., Bowman, H., 2018. Deep temporal models and active inference. Neurosci. Biobehav. Rev. 90, 486–501. https://doi.org/10.1016/J.NEUBIOREV.2018.04.004

Gendlin, E., 1982. Focusing. Bantam Books.

Greenberg, L., 2010. Emotion-Focused Therapy: Theory and Practice. APA Press, Washington, DC.

Gross, J., 1998a. Antecedent- and Response-Focused Emotion Regulation: Divergent Consequences for Experience, Expression, and Physiology. J. Pers. Soc. Psychol. 74, 224–237.

Gross, J., 1998b. The emerging field of emotion regulation: An integrative review. Rev. Gen. Psychol. 2, 271–299.

Gross, J., Levenson, R., 1997. Hiding feelings: The acute effects of inhibiting negative and positive emotion. J. Abnorm. Psychol. 106, 95–103. https://doi.org/10.1037//0021-843X.106.1.95

Hayes, S., Smith, S., 2005. Get Out of Your Mind and Into Your Life: The New Acceptance and Commitment Therapy. New Harbinger Publications, Oakland, CA.

Jack, M.S., Heimberg, R.G., Mennin, D.S., 1999. Situational panic attacks: Impact on distress and impairment among patients with social phobia. Depress. Anxiety 10, 112–118. https://doi.org/10.1002/(SICI)1520-6394(1999)10:3<112::AID-DA4>3.0.CO;2-U

Joffily, M., Coricelli, G., 2013. Emotional Valence and the Free-Energy Principle. PLoS Comput. Biol. 9, e1003094. https://doi.org/10.1371/journal.pcbi.1003094

Kashdan, T., Barrett, L., McKnight, P., 2015. Unpacking Emotion Differentiation: Transforming Unpleasant Experience by Perceiving Distinctions in Negativity. Curr. Dir. Psychol. Sci. 24, 10–16. https://doi.org/10.1177/0963721414550708

Kashdan, T., Farmer, A., 2014. Differentiating emotions across contexts: comparing adults with and without social anxiety disorder using random, social interaction, and daily experience sampling. Emotion 14, 629–38. https://doi.org/10.1037/a0035796

Kavanagh, P., Kahl, B., 2018. Are Expectations the Missing Link between Life History Strategies and Psychopathology? Front. Psychol. 9, 89. https://doi.org/10.3389/fpsyg.2018.00089

Kircanski, K., Lieberman, M., Craske, M., 2012. Feelings into words: contributions of language to exposure therapy. Psychol. Sci. 23, 1086–91. https://doi.org/10.1177/0956797612443830

Lackner, J., 2005. Is IBS a problem of emotion dysregulation? Testing the levels of emotional awareness model, in: Presented at the Annual Meeting of the American Psychosomatic Society.

Lane, R., Anderson, F., Smith, R., 2018. Biased Competition Favoring Physical Over Emotional Pain: A Possible Explanation for the Link Between Early Adversity and Chronic Pain. Psychosom. Med. 80, 880–890. https://doi.org/10.1097/PSY.0000000000000640

Lane, R., Quinlan, D., Schwartz, G., Walker, P., Zeitlin, S., 1990. The Levels of Emotional Awareness Scale: a cognitive-developmental measure of emotion. J. Pers. Assess. 55, 124–34. https://doi.org/10.1080/00223891.1990.9674052

Lane, R., Schwartz, G., 1987. Levels of emotional awareness: a cognitive-developmental theory and its application to psychopathology. Am. J. Psychiatry 144, 133–143.

Lane, R., Sechrest, L., Reidel, R., Weldon, V., Kaszniak, A., Schwartz, G., 1996. Impaired verbal and nonverbal emotion recognition in alexithymia. Psychosom. Med. 58, 203–10.

Lane, R., Sechrest, L., Riedel, R., Shapiro, D., Kaszniak, A., 2000. Pervasive emotion recognition deficit common to alexithymia and the repressive coping style. Psychosom. Med. 62, 492–501.

Lane, R., Weihs, K., Herring, A., Hishaw, A., Smith, R., 2015. Affective agnosia: Expansion of the alexithymia construct and a new opportunity to integrate and extend Freud’s legacy. Neurosci. Biobehav. Rev. 55, 594–611. https://doi.org/10.1016/j.neubiorev.2015.06.007

Lang, P., Bradley, M., Cuthbert, B., 2008. International affective picture system (IAPS): Affective ratings of pictures and instruction manual. Technical Report A-8. University of Florida, Gainesville, FL.

Levine, D., Marziali, E., Hood, J., 1997. Emotion processing in borderline personality disorders. J. Nerv. Ment. Dis. 185, 240–246.

Limanowski, J., Friston, K., 2018. “Seeing the Dark”: Grounding Phenomenal Transparency and Opacity in Precision Estimation for Active Inference. Front. Psychol. 9, 643. https://doi.org/10.3389/fpsyg.2018.00643

Lumley, M., Schubiner, H., Lockhart, N., Kidwell, K., Harte, S., Clauw, D., Williams, D., 2017. Emotional awareness and expression therapy, cognitive behavioral therapy, and education for fibromyalgia: a cluster-randomized controlled trial. Pain 158, 2354–2363. https://doi.org/10.1097/j.pain.0000000000001036

Mayer, J., Salovey, P., Caruso, D., Sitarenios, G., 2003. Measuring emotional intelligence with the MSCEIT V2.0. Emotion 3, 97–105. https://doi.org/10.1037/1528-3542.3.1.97

Melby-Lervåg, M., Hulme, C., 2013. Is working memory training effective? A meta-analytic review. Dev. Psychol. 49, 270–291. https://doi.org/10.1037/a0028228

Mennin, D., Fresco, D., 2014. Emotion regulation therapy, in: Handbook of Emotion Regulation. pp. 469–490.

Metzinger, T., 2017. The Problem of Mental Action, in: Metzinger, T., Wiese., W. (Eds.), Philosophy and Predicitive Processing.

Moeller, S., Konova, A., Parvaz, M., Tomasi, D., Lane, R., Fort, C., Goldstein, R., RZ, G., 2014. Functional, Structural, and Emotional Correlates of Impaired Insight in Cocaine Addiction. JAMA Psychiatry 71, 61. https://doi.org/10.1001/jamapsychiatry.2013.2833

Moors, A., Ellsworth, P.C., Scherer, K., Frijda, N., 2013. Appraisal Theories of Emotion: State of the Art and Future Development. Emot. Rev. 5, 119–124. https://doi.org/10.1177/1754073912468165

Morrish, L., Chin, T.C., Rickard, N., Sigley-Taylor, P., Vella-Brodrick, D., 2019. The role of physiological and subjective measures of emotion regulation in predicting adolescent wellbeing. Int. J. Wellbeing 9. https://doi.org/10.5502/IJW.V9I2.730

Moutoussis, M., Shahar, N., Hauser, T., Dolan, R., 2017. Computation in Psychotherapy, or How Computational Psychiatry Can Aid Learning-Based Psychological Therapies. Comput. Psychiatry 1–21. https://doi.org/10.1162/CPSY_a_00014

Mueller, J., Alpers, G.W., 2006. Two facets of being bothered by bodily sensations: anxiety sensitivity and alexithymia in psychosomatic patients. Compr. Psychiatry 47, 489–495. https://doi.org/10.1016/J.COMPPSYCH.2006.03.001

Ogrodniczuk, J., Piper, W., Joyce, A., 2011. Effect of alexithymia on the process and outcome of psychotherapy: a programmatic review. Psychiatry Res. 190, 43–48.

Owens, A.P., Allen, M., Ondobaka, S., Friston, K.J., 2018. Interoceptive inference: From computational neuroscience to clinic. Neurosci. Biobehav. Rev. 90, 174–183. https://doi.org/10.1016/J.NEUBIOREV.2018.04.017

Panksepp, J., Lane, R., Solms, M., Smith, R., 2017. Reconciling cognitive and affective neuroscience perspectives on the brain basis of emotional experience. Neurosci. Biobehav. Rev. 76, part B, 187–215. https://doi.org/10.1016/j.neubiorev.2016.09.010

Parker, J., Taylor, G., Bagby, R., 2003. The 20-Item Toronto Alexithymia Scale: III. Reliability and factorial validity in a community population. J. Psychosom. Res. 55, 269–275. https://doi.org/10.1016/S0022-3999(02)00578-0

Parr, T., Friston, K., 2018. The Anatomy of Inference: Generative Models and Brain Structure. Front. Comput. Neurosci. 12, 90. https://doi.org/10.3389/fncom.2018.00090

Parr, T., Friston, K., 2017a. Uncertainty, epistemics and active inference. J. R. Soc. Interface 14. https://doi.org/10.1098/rsif.2017.0376

Parr, T., Friston, K., 2017b. Working memory, attention, and salience in active inference. Sci. Rep. 7, 14678. https://doi.org/10.1038/s41598-017-15249-0

Parr, T., Markovic, D., Kiebel, S., Friston, K., 2019. Neuronal message passing using Mean-field, Bethe, and Marginal approximations. Sci. Rep. 9, 1889. https://doi.org/10.1038/s41598-018-38246-3

Peters, A., McEwen, B.S., Friston, K., 2017. Uncertainty and stress: Why it causes diseases and how it is mastered by the brain. Prog. Neurobiol. 156, 164–188. https://doi.org/10.1016/J.PNEUROBIO.2017.05.004

Petrides, K., Mikolajczak, M., Mavroveli, S., Sanchez-Ruiz, M.-J., Furnham, A., Pérez-González, J.-C., 2016. Developments in Trait Emotional Intelligence Research. Emot. Rev. 8, 335–341. https://doi.org/10.1177/1754073916650493

Posner, J., Russell, J.A., Peterson, B.S., 2005. The circumplex model of affect: An integrative approach to affective neuroscience, cognitive development, and psychopathology. Dev. … 715–734.

Potter, C.M., Wong, J., Heimberg, R.G., Blanco, C., Liu, S.-M., Wang, S., Schneier, F.R., 2014. Situational panic attacks in social anxiety disorder. J. Affect. Disord. 167, 1–7. https://doi.org/10.1016/j.jad.2014.05.044

Rottschy, C., Langner, R., Dogan, I., Reetz, K., Laird, A.R., Schulz, J.B., Fox, P.T., Eickhoff, S.B., 2012. Modelling neural correlates of working memory: a coordinate-based meta-analysis. Neuroimage 60, 830–46. https://doi.org/10.1016/j.neuroimage.2011.11.050

Scherer, K., 2009. The dynamic architecture of emotion: Evidence for the component process model. Cogn. Emot. 23, 1307–1351. https://doi.org/10.1080/02699930902928969

Scherer, K., Brosch, T., 2009. Culture-specific appraisal biases contribute to emotion dispositions. Eur. J. Pers. 23, 265–288. https://doi.org/10.1002/per.714

Schnuerch, R., Kreitz, C., Gibbons, H., Memmert, D., 2016. Not quite so blind: Semantic processing despite inattentional blindness. J. Exp. Psychol. Hum. Percept. Perform. 42, 459–63. https://doi.org/10.1037/xhp0000205

Seeley, W., Menon, V., Schatzberg, A., Keller, J., Glover, G., Kenna, H., Reiss, A., Greicius, M., 2007. Dissociable intrinsic connectivity networks for salience processing and executive control. J. Neurosci. 27, 2349–56. https://doi.org/10.1523/JNEUROSCI.5587-06.2007

Segal, Z., Teasdale, J., Williams, J., 2004. Mindfulness-Based Cognitive Therapy: Theoretical Rationale and Empirical Status., in: Hayes, S., Follette, V., Linehan, M. (Eds.), Mindfulness and Acceptance: Expanding the Cognitive-Behavioral Tradition. Guilford Press, New York, NY, NY, pp. 45–65.

Seth, A., 2013. Interoceptive inference, emotion, and the embodied self. Trends Cogn. Sci. 17, 565–573. https://doi.org/10.1016/j.tics.2013.09.007

Seth, A., Friston, K., 2016. Active interoceptive inference and the emotional brain. Philos. Trans. R. Soc. London B Biol. Sci. 371.

Siemer, M., Mauss, I., Gross, J., 2007. Same situation--Different emotions: How appraisals shape our emotions. Emotion 7, 592–600. https://doi.org/10.1037/1528-3542.7.3.592

Smith, R., 2019. The three-process model of implicit and explicit emotion, in: Lane, R., Nadel, L. (Eds.), Neuroscience of Enduring Change: Implications for Psychotherapy. Oxford University Press, p. (in press).

Smith, R., 2017. A neuro-cognitive defense of the unified self. Conscious. Cogn. 48, 21–39. https://doi.org/10.1016/j.concog.2016.10.007

Smith, R., 2016. The relationship between consciousness, understanding, and rationality. Philos. Psychol. 29, 1–15. https://doi.org/10.1080/09515089.2016.1172700

Smith, R, Akozei, A., Killgore, W., Lane, R., 2017a. Nested Positive Feedback Loops in the Maintenance of Major Depression: An integration and extension of previous models. Brain. Behav. Immun. (In Press). https://doi.org/10.1016/j.bbi.2017.09.011

Smith, R., Alkozei, A., Killgore, W., 2019a. Parameters as Trait Indicators: Exploring a Complementary Neurocomputational Approach to Conceptualizing and Measuring Trait Differences in Emotional Intelligence. Front. Psychol. 10, 848. https://doi.org/10.3389/fpsyg.2019.00848

Smith, R., Braden, B., Chen, K., Ponce, F., Lane, R., Baxter, L., 2015. The neural basis of attaining conscious awareness of sad mood. Brain Imaging Behav. 9, 574–587. https://doi.org/10.1007/s11682-014-9318-8

Smith, R., Kaszniak, A., Katsanis, J., Lane, R., Nielsen, L., 2019b. The importance of identifying underlying process abnormalities in alexithymia: Implications of the three-process model and a single case study illustration. Conscious. Cogn. 68, 33–46. https://doi.org/10.1016/J.CONCOG.2018.12.004

Smith, R., Killgore, W., Lane, R., 2018a. The structure of emotional experience and its relation to trait emotional awareness: A theoretical review. Emotion 18, 670–692. https://doi.org/10.1037/emo0000376

Smith, R., Lane, R., 2016. Unconscious emotion: A cognitive neuroscientific perspective. Neurosci. Biobehav. Rev. 69, 216–238. https://doi.org/10.1016/j.neubiorev.2016.08.013

Smith, R., Lane, R., 2015. The neural basis of one’s own conscious and unconscious emotional states. Neurosci. Biobehav. Rev. 57, 1–29. https://doi.org/10.1016/j.neubiorev.2015.08.003

Smith, R., Lane, R., Alkozei, A., Bao, J., Smith, C., Sanova, A., Nettles, M., Killgore, W., 2018b. The role of medial prefrontal cortex in the working memory maintenance of one’s own emotional responses. Sci. Rep. 8. https://doi.org/10.1038/s41598-018-21896-8

Smith, R, Lane, R., Alkozei, A., Bao, J., Smith, C., Sanova, A., Nettles, M., Killgore, W., 2017b. Maintaining the feelings of others in working memory is associated with activation of the left anterior insula and left frontal-parietal control network. Soc. Cogn. Affect. Neurosci. 12, 848–860. https://doi.org/10.1093/scan/nsx011

Smith, R., Lane, R., Nadel, L., Moutoussis, M., 2019c. A computational neuroscience perspective on the change process in psychotherapy, in: Lane, R., Nadel, L. (Eds.), Neuroscience of Enduring Change: Implications for Psychotherapy. Oxford University press, p. (in press).

Smith, R., Lane, R., Sanova, A., Alkozei, A., Smith, C., Killgore, W.W.D., 2018c. Common and Unique Neural Systems Underlying the Working Memory Maintenance of Emotional vs. Bodily Reactions to Affective Stimuli: The Moderating Role of Trait Emotional Awareness. Front. Hum. Neurosci. 12, 370. https://doi.org/10.3389/fnhum.2018.00370

Smith, R., Parr, T., Friston, K.J., 2019d. Simulating emotions: An active inference model of emotional state inference and emotion concept learning. bioRxiv 640813. https://doi.org/10.1101/640813

Smith, R., Quinlan, D., Schwartz, G.E., Sanova, A., Alkozei, A., Lane, R.D., 2019e. Developmental Contributions to Emotional Awareness. J. Pers. Assess. 101, 150–158. https://doi.org/10.1080/00223891.2017.1411917

Smith, R., Sanova, A., Alkozei, A., Lane, R., Killgore, W., 2018d. Higher levels of trait emotional awareness are associated with more efficient global information integration throughout the brain: a graph-theoretic analysis of resting state functional connectivity. Soc. Cogn. Affect. Neurosci. 13, 665–675. https://doi.org/10.1093/scan/nsy047

Smith, R, Thayer, J., Khalsa, S., Lane, R., 2017c. The hierarchical basis of neurovisceral integration. Neurosci. Biobehav. Rev. 75, 274–296. https://doi.org/10.1016/j.neubiorev.2017.02.003

Spaapen, D.L., Waters, F., Brummer, L., Stopa, L., Bucks, R.S., 2014. The Emotion Regulation Questionnaire: Validation of the ERQ-9 in two community samples. Psychol. Assess. 26, 46–54. https://doi.org/10.1037/a0034474

Subic-Wrana, A., Beetz, M., Paulussen, J., Wiltnik, J., Beutel, M., 2007. Relations between attachment, childhood trauma, and emotional awareness in psychosomatic inpatients. Budapest, Hungary.

Subic-Wrana, C., Bruder, S., Thomas, W., Lane, R., Köhle, K., 2005. Emotional awareness deficits in inpatients of a psychosomatic ward: a comparison of two different measures of alexithymia. Psychosom. Med. 67, 483–489.

Swales, Heidi L. Heard, J. Mark G., M., 2000. Linehan’s Dialectical Behaviour Therapy (DBT) for borderline personality disorder: Overview and adaptation. J. Ment. Heal. 9, 7–23. https://doi.org/10.1080/09638230016921

Teigen, K., 1994. Yerkes-Dodson: A law for all seasons. Theory Psychol. 4, 525–547.

Toplak, M., West, R., Stanovich, K., 2014. Assessing miserly information processing: An expansion of the Cognitive Reflection Test. Think. Reason. 20, 147–168. https://doi.org/10.1080/13546783.2013.844729

Toplak, M.E., West, R.F., Stanovich, K.E., 2014. Rational thinking and cognitive sophistication: Development, cognitive abilities, and thinking dispositions. Dev. Psychol. 50, 1037–1048. https://doi.org/10.1037/a0034910

van der Velden, A., Kuyken, W., Wattar, U., Crane, C., Pallesen, K., Dahlgaard, J., Fjorback, L., Piet, J., 2015. A systematic review of mechanisms of change in mindfulness-based cognitive therapy in the treatment of recurrent major depressive disorder. Clin. Psychol. Rev. 37, 26–39. https://doi.org/10.1016/j.cpr.2015.02.001

Versluis, A., Verkuil, B., Lane, R.D., Hagemann, D., Thayer, J.F., Brosschot, J.F., 2018. Ecological momentary assessment of emotional awareness: Preliminary evaluation of psychometric properties. Curr. Psychol. 1–9. https://doi.org/10.1007/s12144-018-0074-6

Waugh, C., Lemus, M., Gotlib, I., 2014. The role of the medial frontal cortex in the maintenance of emotional states. Soc. Cogn. Affect. Neurosci. 9, 2001–9. https://doi.org/10.1093/scan/nsu011

Widen, S., Russell, J., 2008. Children acquire emotion categories gradually. Cogn. Dev. 23, 291–312. https://doi.org/10.1016/j.cogdev.2008.01.002

Wright, R., Riedel, R., Sechrest, L., Lane, R., Smith, R., 2017. Sex differences in emotion recognition ability: The mediating role of trait emotional awareness. Motiv. Emot. https://doi.org/10.1007/s11031-017-9648-0

Xin, F., Lei, X., 2015. Competition between frontoparietal control and default networks supports social working memory and empathy. Soc. Cogn. Affect. Neurosci. 10, 1144–1152. https://doi.org/10.1093/scan/nsu160

Zemack-Rugar, Y., Bettman, J., Fitzsimons, G., 2007. The effects of nonconsciously priming emotion concepts on behavior. J. Pers. Soc. Psychol. 93, 927–939. https://doi.org/10.1037/0022-3514.93.6.927

